# Consolidated sustainable organic nanozyme integrated with Point-Of-Use sensing platform for dual agricultural and biological molecule detection

**DOI:** 10.1101/2024.10.02.616341

**Authors:** Dong Hoon Lee, Mohammed Kamruzzaman

## Abstract

The newly introduced organic nanozymes offer a promising alternative to inorganic nanozymes by sustainable and cost-effective solutions. Given the potential of nanozyme-based Point-of-Care (PoC) systems for on-demand molecule sensing; these user-friendly Point-of-Use(PoU) platforms can be highly beneficial for real-world agricultural and biological applications. Herein, a novel, fully organic compound/monomer-based, sustainable organic nanozyme is introduced, that exhibits peroxidase-like enzyme-like catalytic activity. Utilizing a unique self-assembled particle fabrication process, homogenous nanozyme are fabricated within a short time (within 50 nm, *D_90_*), exhibiting decent kinetic profiles (*K_m_*= 0.046 mM, H_2_O_2_), and degradability for improved waste management after its use. OM nanozyme-incorporated colorimetric sensing platforms successfully detect popular agricultural toxic biomolecules and biological molecules while achieving decent molecule sensing performance (practical LOD, glyphosate: at least 1pgmL^-1^, glucose: 3.47 ngmL^-1^) with high analytical sensitivity and selectivity. Additionally, a smartphone image-based polynomial regression analysis system and paper-microfluidic hardware were developed to facilitate PoU applications in a lab-free environment with satisfactory analytic sensitivity (at least 1.6μgmL^-1,^ for glyphosate, and glucose). This sustainable OM nanozyme, coupled with the PoU platform, is envisioned to have broad applications in agricultural and biological systems.

## 1. Introduction

Nanozyme [1–5] are promising nanomaterials that have demonstrated enzyme-like catalytic performance with higher stability, offering a potential alternative to natural enzymes for diverse applications. Most nanozymes are made from inorganic materials, however, recent studies have introduced organic-material-based nanozyme, known as ‘organic nanozymes’[6]. These organic nanozymes partially overcome the chronic problems associated with inorganic nanozymes [6], such as toxicity, and high costs, offering more sustainable properties. Organic nanozymes are made of relatively non-toxic materials (e.g., *LD _50_*> 5,000, referred to as ‘practically non-toxic’), and their unit price is lower than the inorganic materials (e.g., gold, 1g = 400 USD, approximately)[6]. Despite their potential, organic nanozymes are still in the early stages of development, and many of their constituent materials contain a significant inorganic portion [6]. The advent of nano hydrogel nanozyme [7] and OC (organic-compound-based, mostly for agricultural and eco-friendly) nanozyme [8–10] partially resolve these limitations: These nanozymes do not rely on ‘already made’ inorganic nanostructures or catalysts to use them as an ‘active sites’ but contain transitional metal ion-based ligands as active sites which generated in the middle of the nanoparticle formation process [7–10].

Particularly, previous OC nanozymes offer a complete one-pot, partially cofactor-mimetic, homogenous nanozyme generation method without the need for chemical engineering process, heat treatment, or ionic gelation, which contributes to higher particle generation yield [8–10]. Although, these nanozymes exhibit suitable morphological structures, they typically require a polymer-based particle stabilizer to maintain their spherical framework. However, the use of higher molecular weight polymers can lead to larger nanozyme dimensions. Thus, it is necessary to fabricate nanozymes without a polymer-stabilizer utilizing a double-chelation method with organic compounds and monomers. This **‘**pseudo-amalgamation**’** approach creates a stable nanoscale framework and facilitates degradation after use due to the cleavage of the coordinated bond at the designated condition. Overall, this method provides smaller organic nanozymes that can deform, and degrade under designated conditions, highlighting their sustainable outlook.

Besides, various PoC applications using nanozymes are being consistently developed [11]. These nanozyme-based PoC platforms provide great potential for user-focused molecule diagnosis in lab-free environments. However, the limitations of inorganic nanozyme, mainly their high costs and toxicity, used PoC platform pose significant challenges for real-world implementation. The use of organic, and sustainable nanozyme made from biologically non-toxic materials (e.g., evaluated by *LD_50_* values) and having degradable properties offer a promising alternative for sustainable, real-world applications. Thus, organic nanozyme-based PoC applications hold a substantial practical perspective.

Detecting toxic agrichemical, alongside biological molecules sensing, is crucial for agricultural safety. A PoU application capable of monitoring both agricultural toxic molecules and biomolecules simultaneously can be beneficial. In this paper, a non-particle stabilizing polymer containing next-generation organic nanozyme (OM nanozyme) is proposed, which possesses peroxidase-like enzyme-like catalytic activity. The modified organic-compound/monomer based nanozyme fabrication strategies, via double-chelation methods, have successfully generated small and homogenous nanozymes (mostly within 50 nm of physical diameter) containing partially mimicked cofactor mimetics (e.g., Fe_x_-N_y_, Fe_x_-O_y_ sites, etc). The OM nanozyme possesses decent substrate affinity calculated by Michael Menten’s kinetic analysis (e.g., *K_m_* = 0.046 mM, H_2_O_2_), and particle deformation (related to degradability) at moderate temperature (e.g., 46 °C) was confirmed for its sustainable outlook. The OM nanozyme incorporated customized colorimetric sensing platform enables the fast (up to five minutes) detection of agricultural herbicide (e.g., glyphosate) and biological molecule (e.g., glucose), with having decent analytic sensitivity and selectivity, with mixture selectivity (similar to specificity). Furthermore, a novel smartphone camera-based readout system, using with regression analysis of HSV values, replaces the need for t conventional UV-VIS spectroscopical readouts. Additionally, a microfluidic paper-based analytic device (µPADS) was developed to facilitate PoU applications. This OM nanozyme and incorporated PoU system highlights the diversity of organic nanozymes and contribute to effective molecule sensing for applications in agricultural safety and human health.

## 2. Materials and methods

### 2.1 Materials

Urea from Thermo Scientific, Acrylamide(A8887), Iron(ll) sulfate heptahydrate (F8633), Sodium Acetate (236500), ABTS (2,2′-Azino-bis (3-ethylbenzothiazoline-6-sulfonic acid) diammonium salt)(A1888), TMB(3,3,5,5’-Tetramethylbenzidine), and OPD(o-Phenylenediamine)(P23938)(860336), Glucose oxidase from aspergillus, Glyphosate (45521), Whatman Cellulose Grade 1 chromatography paper(WHA3001861) from Sigma-Aldrich were used.

### 2.2 Fabrication of Organic compound/monomer-based nanozyme (OM nanozyme)

Iron sulfate heptahydrate powder was dissolved in deionized water and serial dilutions were performed to make the target solution (final concentration of 1mg mL^-1^). Urea powder was similarly prepared and serial diluted to reach the target concentration (final concentration of 1mg mL^-1^). These two solutions were then mixed in a 1:1 volume ratio and reacted to form the initial coordinated bond under the stirring condition at room temperature (around 25°C). Acrylamide solution (final concentration 1mg mL^-1^) was added dropwise while stirring. OM nanozymes were collected through centrifugation (10 minutes at a maximum of 4000 rpm). The sample was washed several times (if necessary) using deionized water for further analysis.

### 2.3 Morphological analysis of OM nanozyme

Morphological analyses were conducted using transmission electron microscopy (TEM) and scanning electron microscopy (SEM). Specifically, TEM images were acquired with a Tecnai G2 F20 S-Twin instrument from ThermoFisher Scientific, operating at a maximum of 200 kV. SEM images and Energy-dispersive X-ray spectroscopy (EDS) profile were obtained using a Hitachi S-4800 SEM, operating at an accelerating voltage of up to 100 kV. Nanoparticle tracking analysis (NTA) profiles were procured using a Malvern Panalytic Nanosight NS300 equipped with an sCMOS camera and a blue488 laser. Dynamic light scattering (DLS) and zeta potential profiles were acquired using a Malvern Zetasizer from Malvern Panalytic. Additional spectroscopical characterization related to the morphological properties was conducted via a Biotek Gen 5 microplate reader.

### 2.4 Chemical bonding and related characterization of OM nanozyme

XPS X-ray photoelectron spectroscopy (XPS) and Powder X-ray crystallography(pXRD) analyses were employed to verify the chemical bonding. The XPS measurements were acquired using a Kratos Axis Supra plus instrument, with dual anode Al Ka/Ag La X-ray source serving as the excitation source (sample substrate: silicon wafer). The pXRD measurements were conducted using Cu Kα x-ray radiation, scanning two thetas from 5° to 100° angle) (sample substrate: silicon wafer).

### 2.5 Sample preparation of the EPR experiment

The Electron paramagnetic resonance (EPR) profile was obtained from the EMXplus EPR instrument (Bruker, USA). The OM nanozyme (default concentration, 2.72±0.74 ×10^−13^M), H_2_O_2_ (1mM, final concentration), sodium acetate buffer (pH4, 0.1M), and DMPO dissolved in sodium acetate buffer (concentration up to 0.5mgmL^-1^) was used for the sample preparation.

### 2.6 Optimization and characterization of experimental condition for colorimetric sensing application

Optimization of the OM nanozyme activity based on pH, and nanozyme concentration was conducted, as described in previous studies [8–10]. Room temperature (25 °C) and pH 4 were selected for the entire experiment as these conditions resulted the maximum absorbance endpoint.

### 2.7 Peroxidase-like activity validation via colorimetric assay with OM nanozyme

Peroxidase activity was evaluated using colorimetric assays with coloring substrates such as ABTS, TMB, and OPD). A 96-well plate was prepared with 15 μL of OM nanozyme (default concentration:2.72±0.74 ×10^−13^M), 15 μL of H_2_O_2_ (final concentration of 100 μM), and 60 mM of ABTS, 8mM of TMB, 5mM of OPD in a 105 μL solution of sodium acetate-acetic acid buffer [final concentration: 70 mM pH 4.0]. Concurrently, control groups were established by substituting H_2_O_2_ with deionized water and replacing the nanozyme with buffer. The catalytic oxidation of ABTS, TMB, or OPD was assessed by monitoring the absorbance changes of their oxidized form at λmax = 417 nm, 652nm, and 450nm, respectively. Absorbance spectra were obtained over a wavelength range of 400 to 500 nm (for TMB, 500-800nm), with each group scanned three times.

### 2.8 Steady-state kinetic analysis of OM nanozyme

Steady-state kinetic measurements were conducted in kinetic mode utilizing Varian Cary 5G and Agilent Cary 5000 instruments. The reaction mixture comprised 700 μL of NaAc buffer, 100 μL of the OM nanozyme (default concentration), 100 μL of the substrate (e.g., H_2_O_2_) in a transparent cuvette, along with varying concentrations of TMB. Absorbance endpoint peak signals were consistently recorded for 1 minute. Kinetic parameters were derived from the Michaelis-Menten equation, with Michaelis constant (*K_m_*) and maximum initial velocity (*V_max_*) obtained from Lineweaver-Burk plots and the corresponding Michaelis-Menten equation, described as ν = *V_max_* × [S]/ (*K_m_* + [S]), where ν denotes the initial velocity and [S] represents the substrate concentration.

### 2.9 Glyphosate detection via OM nanozyme-based colorimetric sensing platform

Glyphosate was selected as the target agricultural herbicide. It was dissolved in deionized water and conducted serial dilutions to create various concentrations. This solution was then incorporated with a previously established colorimetric assay (default concentration of OM nanozyme + 0.1M sodium acetate buffer at pH 4 + 1mM H_2_O_2_). Glyphosate solution(10% of the total volume) was added to the column and thoroughly mixed with a pipette. ABTS (6mM final concentration) was used as the target coloring substrate. Endpoint absorbance measurements were conducted within 3 minutes of assembling all components in the 96-well plate. Aanalytic selectivity and mixture selectivity (similar to specificity) experiment was conducted by substituting the glyposate solution with an equal volume of various agricultural and biological molecule solutions (e.g., alanine, glycine, histidine, gluthation, calcium choloride, potassium choloride) to confirm the analytic selectivity. A mixture of chemical solutions (pseudo-real sample) was prepared to mimic the actual agricultural samples, containing high concentration (up to 1mgmL^-1^) of 6 or more agricultural, and biomolecules (e.g., alanine, glycine, histidine, gluthation, calcium choloride, potassium choloride) dissolved in deionized water. Two sample sets were prepared: one containing glyphosate (up to 1mgmL^-1^) and other without glyposate (replaced with the same volume of NaAc Buffer) to confirm the OM nanozyme-based molecule sensing platform’s analytic specificity.

### 2.10 Glucose detection via OM nanozyme-based colorimetric sensing platform

Glucose was selected as the target biological molecule. D-glucose powder was dissolved in deionized water and serial diluted to create various concentrations. Similar to the previous glyphosate detection, this solution was used in the previously established colorimetric assay (default concentration of OM nanozyme + 0.1M sodium acetate buffer at pH 4). Glucose oxidase enzyme (100μgmL^-1^) was dissolved in the deionized water and added (10% of the total volume) after inserting buffer and OM nanozyme. The glucose solution (10% of the total volume) was added to the column and thoroughly mixed with a pipette. For measurement purposes, ABTS (6 mM final concentration) was incorporated. The 96-well plate and the UV-VIS spectroscopy were used (Biotek 5 microplate reader) for absorbance signal measurement. Endpoint absorbance signal measurements were conducted within 5 minutes of assembling all components in the 96-well plate. An assay using a sucrose solution was also conducted to confirm the analytic (sugar) selectivity.

### 2.11 OM nanozyme’s degradability analysis

OM nanozyme dispersed in deionized water was incubated in a temperature-adjustable laboratory incubator (up to 46 °C) for a maximum of 24 hours. OM nanozyme was re-collected through a centrifuge down process (10 minutes, maximum 5000 rpm) and resuspended in deionized water. The sample was subsequently mounted on a silicon wafer for SEM analysis. Four time points (0, 3, 18, 24 hours) were selected for morphological analysis to confirm the degradation. EDS quantification and XRD analysis were carried out, and control and 24-hour samples were designated for comparison.

### 2.12 Preparation and optimization for the smartphone image-based analysis system for colorimetric sensing

The image was directly captured using an iPhone 8 after putting all components for the colorimetric assay. The distance between the well plate (or µPADS) and the camera is around 30cm (Lux value, 230 ∼ 400). The image file was saved as ‘JPEG’ for further image processing. The MATLAB APP program was developed for obtaining the optical values of a certain area, and optical values (e.g., H, S, V) were manually captured through the squared-shape designation from this APP. The extracted values were then sorted and prepared for the regression analysis (e.g., polynomial regression analysis). The goal of regression analysis was to identify the best fitting model with the highest R^2^ value to obtain a similar linearity as UV-VIS spectroscopical methods.

### 2.13 Paper microfluidics chip(μPADS) for conducting PoU, dual molecule sensing application

The μPADS design was carried out using a 3D CAD app for the scratch steps, and the final design was confirmed using Microsoft PowerPoint. Cellulose chromatography paper was used as the substrate, and the pattern was printed using an inkjet printer. The printed chips were then incubated on a hot plate at over 120 °C for up to 30 minutes. The hydrophobic polymer solution (e.g., Polylactic acid) was manually coated with the edge of the well.

## 3. Characterization of OM nanozyme

### 3.1. Morphological and chemical Analysis

The OM nanozyme which is expected to have a Fe-based active site within its matrix was synthesized through a customized double chelation-based particle generation process (**Figure 1 a, SI Figure 1**). The fabrication steps were optimized through a screening process, and the configuration of the OM nanozyme was selected due to its enhanced enzyme-like catalytic performance (**SI Figure 2**). Unlike other iron-based nanoparticles (e.g., Fe_3_O_4_ nanoparticle, etc), the OM nanozyme is transparent, which is suitable for optical sensing applications (**Figure 1 b, SI Figure 3**). Morphological analysis was conducted using electron microscopic techniques, including TEM and SEM with EDS (**Figure 1 c-e**). SEM analysis verified the spherical structure and uniform size of OM nanozyme, demonstrating successful and stable particle structure generation (**Figure 1 c**). The EDS profiles showed the presence of all designated elements (e.g., iron, nitrogen, carbon, and oxygen) in the OM nanozyme’ framework (**Figure 1 d**), while the TEM profile confirmed its spherical morphological properties (**Figure 1 e, SI Figure 4**). The size distribution of OM nanozyme was measured using NTA, and most of the nanozyme’s diameters are within 50 nm. The surface charge in a neutral pH environment is approximately 6mV (**Figure 1 f, SI Figure 5**).

**Figure 1.**
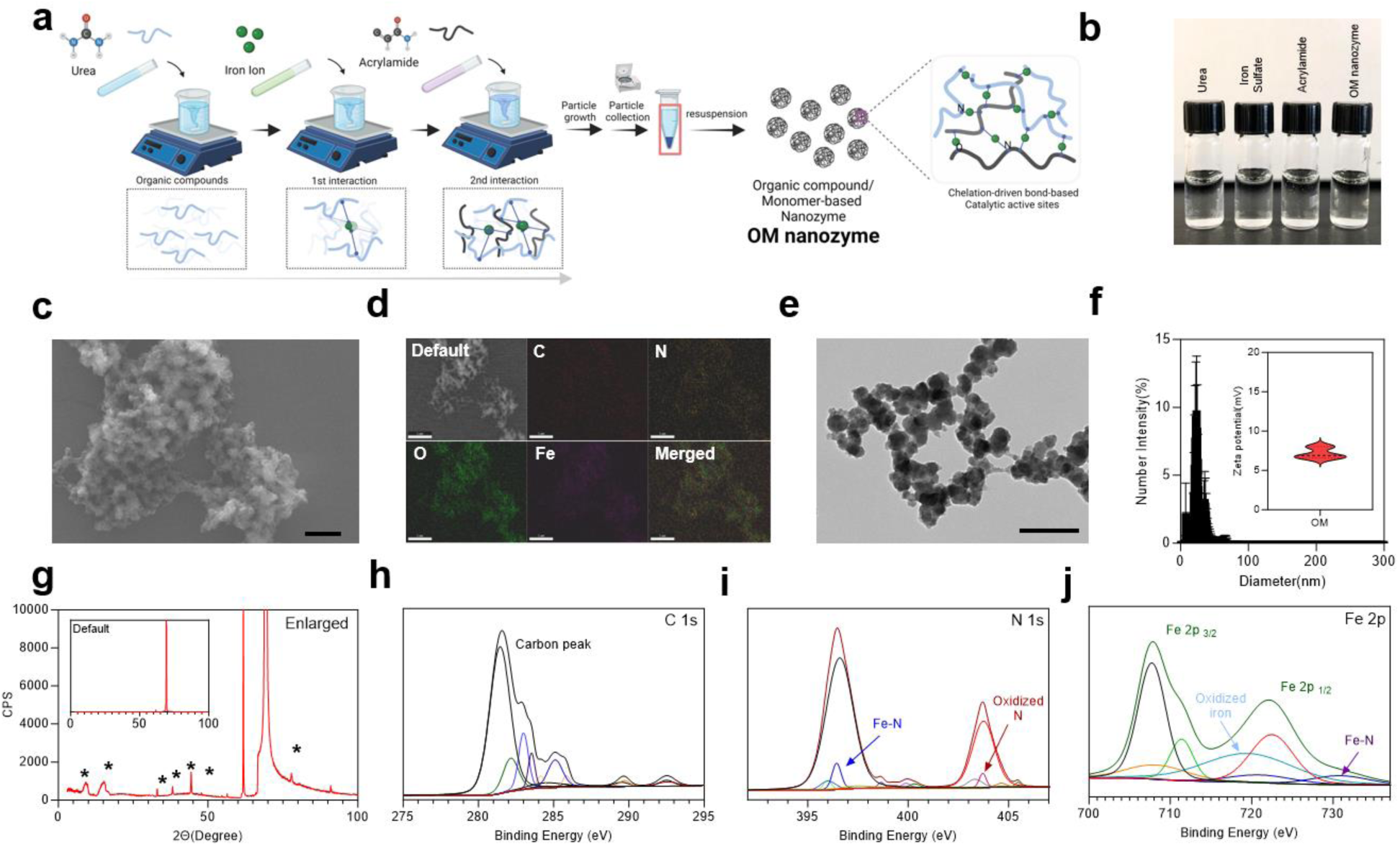
Fabrication and characterization of OM nanozyme. **a.** Illustration of OM nanozyme fabrication via double chelation process. **b.** OM nanozyme and its constituent materials in the solution phase and their transparency. **c.** SEM image of OM nanozyme (Scale = 500nm). **d.** EDS of OM nanozyme(Scale = 1μm). **e.** TEM image of OM nanozyme (Scale = 500nm). **f.** Size distribution and surface charge of OM nanozyme(N=3). **g.** Enlarged XRD analysis graph of OM nanozyme(inserted figure = default scaled graph). **h-j**. XPS deconvolution analysis of OM nanozyme(from left, C 1s, N 1s, Fe 2p).

XRD analysis was carried out to confirm the crystallinity of the OM nanozyme (Figure 1 g)[12–14]. Eight peaks were observed in the pXRD, pattern at 9.08, 15.2, 33.04, 38.16, 44.36, 47.47, 56.38, and 90.92(degree, 2θ) which corresponds to the d-spacing(Å) as approximately 9.40, 5.82, 2.71, 2.36, 2.04, 1.90, 1.63, 1.06, respectively. Since all synthesis steps were conducted at room temperature (around 25°C) and 1 atmospheric pressure, only a limited number of iron-based products can form within the OM nanozyme’s frameworks [12–15]. The peak at 9.08 degrees and 15.2 degrees matches that of hydrate iron oxide, indicating the presence of a basal plane (related to the interlayer spacing). The peak of 33.04 degrees can be attributed to Fe_3_O_4_ (magnetite) with an expected (311) plane [12]. The peak at 38.16 degrees corresponds to Fe_3_O_4_ (400). The peaks 44.36 degrees and 47.47 degrees (200) are associated with Fe_4_N(Iron nitride)[13]. The peak at 56.38 degrees corresponds to either Fe₃N (Iron Nitride) (440) or the (211) plane of Fe_3_N [14]. Lastly, the peak of 90.92 degrees can be assigned to the (533) for Fe₃O₄ or the (311) plane of Fe₄N[15]. To further confirm the chemical bonding information of the OM nanozyme, XPS analysis was conducted (**Figure 1 h-j, SI Figure 6**). The analysis targeted 4 elements and their orbitals : C 1s, O 1s, N 1s, and Fe 2p. For C 1s (**Figure 1 h**), a peak at 294.49 eV indicates the presence of a carbon peak[16], and the *π* - *π* * transition from carbon was observed at 292. 51eV[17]. In the N 1s spectrum (**Figure 1 i**), the binding energy range was confirmed from 393 eV to 407 eV. The peak at 396. 58 eV indicates the Fe-N bond [18], which matches the Fe-N bond observed in the Fe 2P orbital. The peak at 397.46 eV suggests a sp^2^ hybridized N[19], indicating the formation of an imine-like structure due to the Fe-N based bonding procedure with acrylamide, where sp^3^ hybridized N is generally expected [19]. The 403.3 eV indicates oxidized-like N, due to the result of a coordinated bond [20]. For Fe 2p (**Figure 1 j**), the binding energy range was confirmed from 700 eV to 738 eV. The peak at 707.73 eV indicates the Fe^2+^ 2p_3/2_,[21]while the 719.52 eV peak indicates the satellite peak at a higher binding energy value of Fe^2+^ 2p_3/2_. [22]. The peak at 722.5 eV corresponds to Fe 2p_1/2_[23], and the peak at 719.52 also matches Fe^0^ and oxidized iron [24], indicating the presence of iron oxide (Fe-O sites) in the matrix[24], which could serve as active sites. Additionally, the peak at 730.8 eV suggests the presence of Fe-N_x_ bond [25], which may be designated as active sites. Lastly, FT-IR analysis was conducted to cross-check the chemical bonding information of OM nanozyme (**SI figure 7**).The analysis detected key active site bonds, including Fe-N (around 882 cm^-1)^and Fe-O (around 575 cm^-1^) which matches with the XPS data, confirming the presence of these bonds in the OM nanozyme.

### 3.2. Peroxidase-like enzymatic activity of OM nanozyme

The OM nanozyme was designed to partially mimic the cofactor of Horseradish Peroxidase(HRP). Given the presence of active site structures (e.g., Fe_x_-N_y_ or Fe_x_-O_y_, etc) in the OM nanozyme(**Figure 1 h-j**), it is expected to exhibit peroxidase-like (POD) catalytic activity. This suggests that the OM nanozyme can use H_2_O_2_ as an initial substrate, producing hydroxyl radical as an intermediate and generating H_2_O and oxidized substrates as final products (**SI Figure 1**). To understand its catalytic activity, a colorimetric assay with coloring-substrates and EPR analysis were conducted (**Figure 2 a-h**). ABTS, TMB, and OPD were used in the colorimetric assay to assess POD activity (**Figure 2 a**). EPR analysis (**Figure 2 b**) four major peaks were observed with the OM nanozyme+H_2_O_2_+DMPO in the EPR profile, confirming the OM nanozyme selectively consumes H_2_O_2_ as the initial substrate, generating hydroxyl (or hydroxyl-like) radical due as an intermediate reaction step via Fenton-like reaction through the Fe-based cofactors located in the OM nanozyme(**SI Figure 1, Figure 2 b**). The reference group, which contained the OM nanozyme and DMPO but lacked H_2_O_2_, did not show any peaks, confirming the POD activity of the OM nanozyme (**Figure 2 b**). A coloring substrate-based colorimetric assay using OM nanozyme was conducted to verify its POD activity indirectly and utilization for further colorimetric assay (**Figure 2 c-e**). Absorbance scanning was performed in the range of wavelength 400-500 nm for ABTS and OPD, and 500-800 nm for TMB, providing peak values at the λ_max_ of each coloring substrate (e.g., λ_max_ ABTS = 417nm, λ_max_ TMB = 652nm, λ_max_ OPD = 450 nm), demonstrating successful oxidization of those substrates, accelerated by OM nanozyme’s POD activity (**Figure 2 c-e**). The control groups, which contained neither H_2_O_2_ nor the OM nanozyme, did not show significant color change reaching the peak endpoint at the λ_max_ point of the substrates(**Figure 2 c-e**), indicating OM nanozyme is the only element to exhibit enzyme-like catalytic activity and selectively demonstrates POD activity without oxidase-like activity, consistent with the previous EPR analysis (**Figure 2 B, SI Figure 8**). The time-dependent increase in the absorbance signal was measured for up to 400 seconds using the same groups (OM + H_2_O_2_, OM+ H_2_O, Buffer+ H_2_O_2_) as previously used for the colorimetric assay, showing that the OM nanozyme can continuously consume the substrate (e.g., H_2_O_2_) until saturation (**Figure 2 f-h**) [7–9]. Before kinetics analysis, characterization of pH-dependent, concentration-dependent, and substrate-concentration-dependent absorbance signal linearity was measured, resulting in a standard curve (substrate = H_2_O_2_, R^2^ = 0.9868, and substrate concentration/linear range: 0 - 500 µM) (**SI Figure 9**). A steady-state kinetic assay was then carried out to understand the enzyme-like kinetic characteristics of the OM nanozyme (**Figure 2 i-l**). To ensure accurate kinetic analysis using a coloring substrate (e.g., TMB), the initial substrate concentration was lowered. In this case, the H_2_O_2_ concentration was reduced to one-tenth of that used in a previous colorimetric assay to prevent the fast oxidation of the coloring substrates from reaching the saturated point. By obtaining catalytic information, the kinetic profiles and their parameters were calculated using the conventional Michaelis-Menten equation and Lineweaver Burk plot for further analysis of the kinetics behavior of the OM nanozyme. The *K_m_* of OM nanozyme on H_2_O_2_ was 0.046 mM, *V_max_* = 1.522 µMs^-1^, (**Figure 3 F,G, SI Figure 9, SI Table 1**): This profile suggests that the nanozyme binds strongly and efficiently to the substrate at low concentrations. Based on the comparison with the recently developed nanozymes (**SI Table 1**), the OM nanozyme exhibits a decent *k_m_*, indicating higher substrate affinity, which is advantageous for sensing applications.

**Figure 2.**
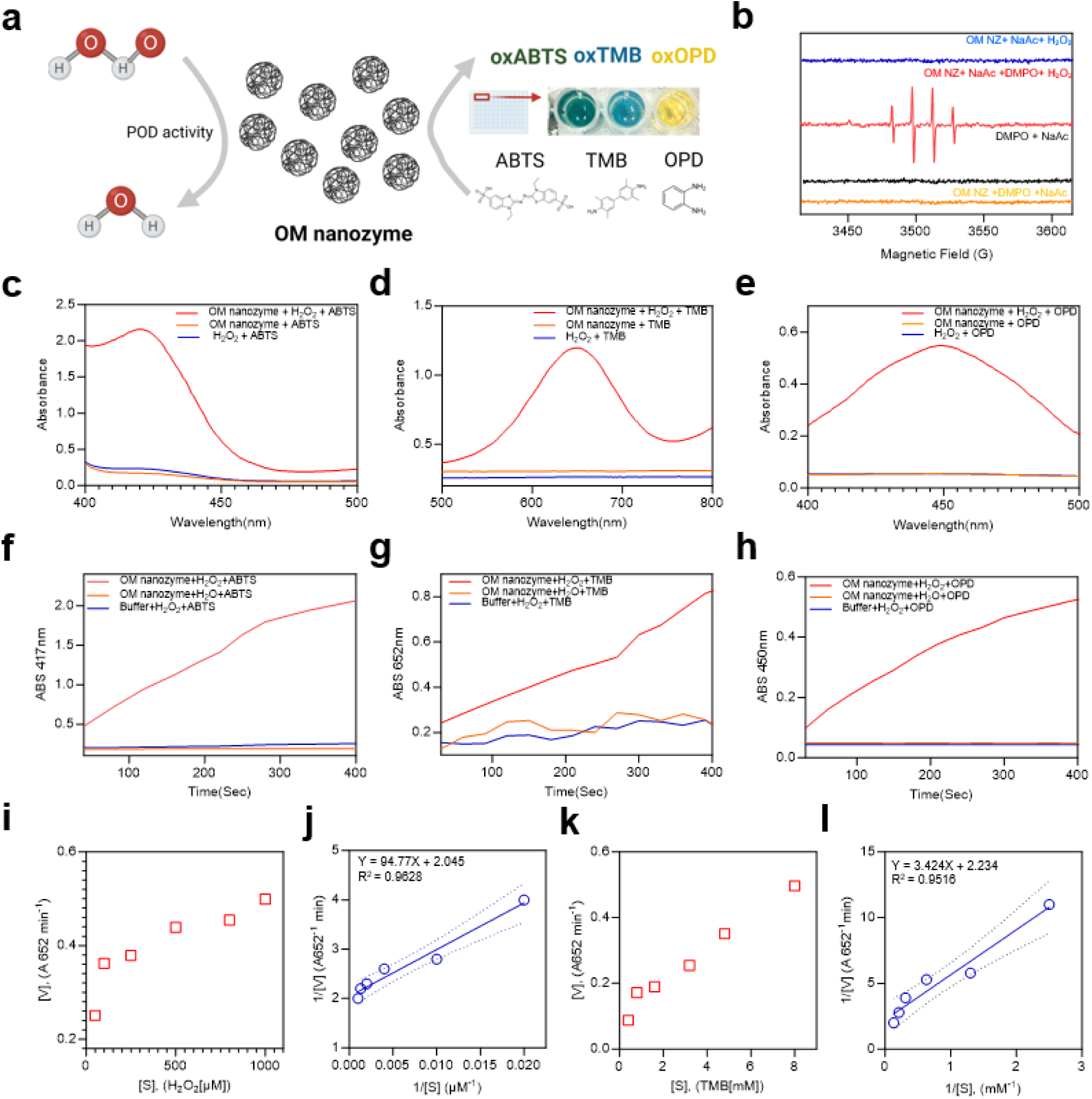
Enzyme-like catalytic activity of OM nanozyme. **a.** Illustration of OM nanozyme-based colorimetric assay. **b.** EPR analysis of OM nanozyme for POD activity verification. **c-e.** OM nanozyme-coloring substrate assay-based, Absorbance spectra measurement via UV-VIS-NIR spectroscopy. (Red: OM nanozyme + H_2_O_2_ + coloring substrates, Orange: OM nanozyme+ deionized water+ Coloring substrates, Blue : Buffer+ H_2_O_2_+ coloring substrates). **f-h**, Time-dependent POD activity measurement. Substrate, **f:** ABTS, **g:** TMB, **h:** OPD. **i-l** Kinetic analysis of OM nanozyme(Left: Curve of the Michael Menton equation and the Right: Lineweaver-Burk plot, **i-j** : H_2_O_2_, **k-i** : TMB).

**Figure 3.**
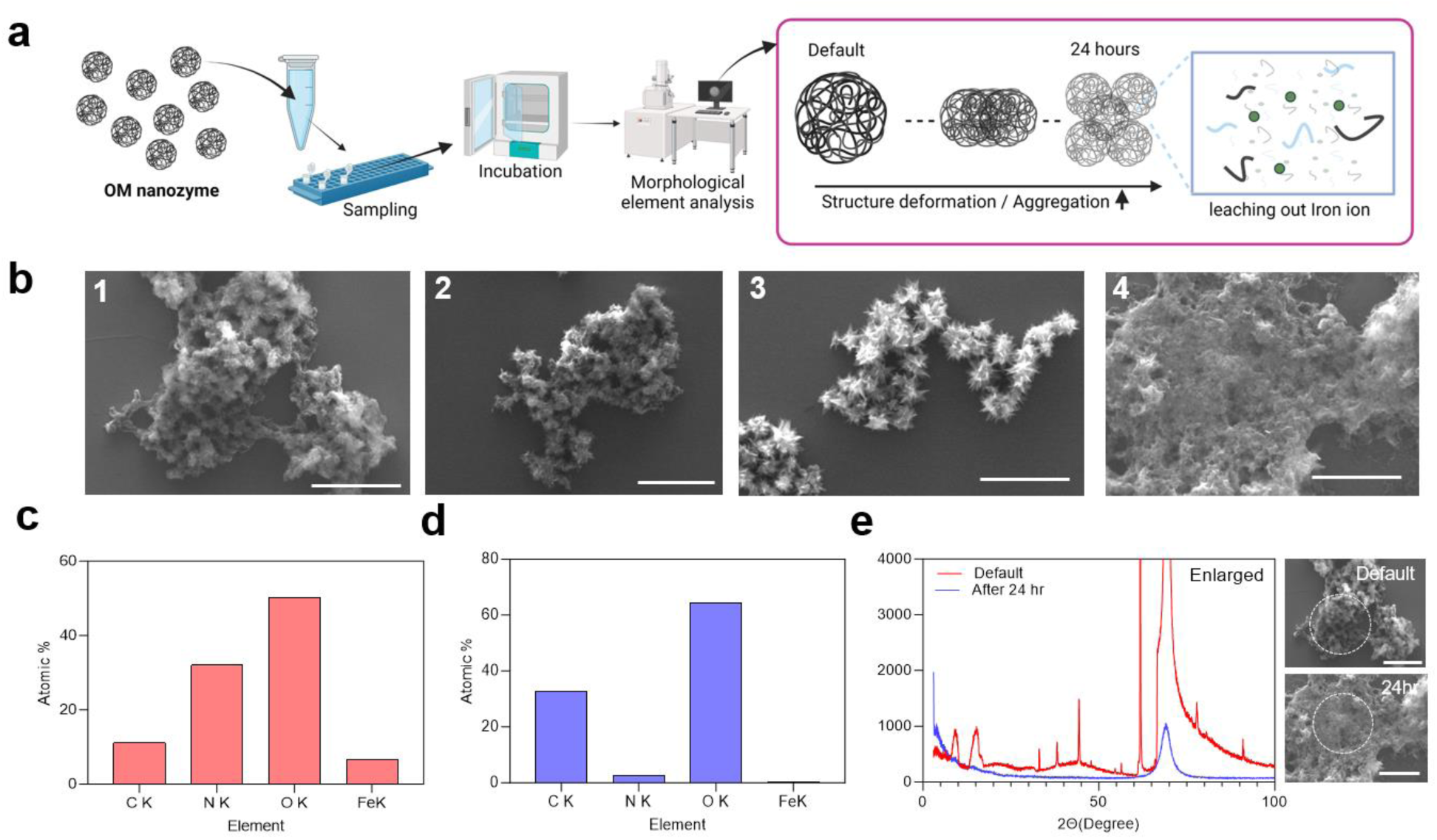
Sustainable perspective of OM nanozyme(degradability). **a.** Illustration of OM nanozyme’s framework degradation in moderate temperature(46 °C), **b.** Incubation time-dependent Representative SEM image(Scale Left from right, 1μm). **c-d**. Element quantification profile comparison from **c:** default to **d:** 24 hours, referred to significant structure deformation. **e.** XRD profile comparison between default and the 24 hr incubation(Right, reference SEM image for the demonstration).

### 3.3 Degradability/deformability of OM nanozyme

Recent research on nanomaterials consistently focuses on sustainability, particularly considering the impact of nanomaterials on the environment after intended use [26]. Most inorganic nanozymes are resistant to degraded under moderate temperatures and pH environments, raising concerns about waste management in real-world applications, similar to the ongoing issue with microplastics. To overcome these drawbacks, the OM nanozyme was designed to deform its physical framework at a moderate temperature, around 46 °C or above. This temperature can be easily achieved in everyday settings (e.g., household water baths). Thus, the assessment of the degradable condition of OM nanozyme is not complicated.

The primary driving force behind the formation of the OM nanozyme is the double chelation-mediated coordinated bond between iron ion and functional groups from organic compounds/monomers. Thus, breaking these coordinated covalent bonds is crucial to achieving particle deformation (**Figure 3 a**). This coordinated bond cleavage happens due to the thermal decomposition when exposed to a certain temperature (e.g., shifting the equilibrium toward a dissociated state/ exothermic formation). To confirm this hypotyposis, both morphological and XRD analyses were conducted to verify the OM nanozyme’s deformation and iron ion or iron-based ligand escape confirmation at a moderate temperature (e.g., 46 °C) over a 24-hour incubation period. SEM analysis provides straightforward information on particle deformation, showing clear morphological changes between the default state and 24-hour incubation (**Figure 3 b**). Additionally, EDS quantification analysis was conducted to compare the elemental atomic ratios before and after 24-hour incubation of OM nanozyme(**Figure 3 c-d**): The iron atomic ratio is a crucial factor indicating deformation, as the cleavage of coordinated bond triggers the release of iron ion from the particle-matrix. The results indicated a drastic decrease in iron content after incubation (e.g., from 6.52 to 0.26, aa%), indirectly confirming the OM nanozyme’s deformation indirectly. Further analysis using XRD showed the disappearance (**Figure 3 e**) of all designated Fe-based composition peaks(e.g., Fe_3_O_4_, Fe_3_N, etc.), in the 24-hour incubation samples matching with the EDS quantification profiles that suggest the escape of iron ion and iron-based ligands. These findings indicate that the OM nanozyme could deform its framework in a relatively moderate condition in a short period, highlighting its potential for a more sustainable environmental impact compared to conventional inorganic nanozyme and related nanoscale substances. In addition to its degradability, the storage stability of the OM nanozyme was confirmed, and the half-life within the solution phase is around 11 days when stored at 4 °C (**SI Figure 10**).

### 3.4. OM nanozyme-based colorimetric sensing platform for dual molecule detection

OM nanozyme-based biomolecules sensing platform was developed for agricultural and biomolecule detection. In agriculture, pesticides are common analytes, with glyphosate being a widely used herbicides, and accurate detection of glyphosate is crucial due to its potential impact on human health. For biological molecule detection, glucose was selected as a target since it has multiple reference sensors with the nanozyme-colorimetric assay.

The customized sensing mechanisms were adapted from the previous study [8]. It was found that glyphosate may interfere with the oxidation of the coloring substrate in the middle of the POD activity as it can shield the active sites/or the framework of the nanozymes (**Figure 4 a**). The OM nanozyme-based sensing system employed this methodology for glyphosate detection [8]. The analytic sensitivity was confirmed by characterizing the concentration-dependent absorbance signal using serially diluted glyphosate solution (**Figure 4 b**), with a practical LOD estimated to be at least 1pgmL^-1^. To assess analytic selectivity, the OM nanozyme-based colorimetric assay was tested with a solution containing various agricultural and biological molecules-(**Figure 4 c**). As glyphosate decreases the absorbance signal, the results confirmed that that the presence of other molecules did not significantly influence the signal reduction. Further statistical analysis (one-way ANOVA) validated this selectivity, showing a highly significant adjusted P value (****<0.0001) for glyphosate compared to positive control. To prove the potential glyphosate detection in agricultural samples, a pseudo-real-sample was prepared by mixing six or more agricultural and biological molecules at high concentrations (up to 1mgmL^-1^)(**Figure 4 d-e, SI Figure 11**). The OM nanozyme-based colorimetric assay was conducted on this pseudo-real-sample., The results indicated a significant detection of glyphosate compared to the glyphosate-negative control, which did not contain glyphosate in the mixture of chemicals (**Figure 4 d**). Therefore, affordable analytic sensitivity, selectivity, and mixture selectivity (in other words, specificity) of glyphosate detection using OM nanozyme was verified.

**Figure 4.**
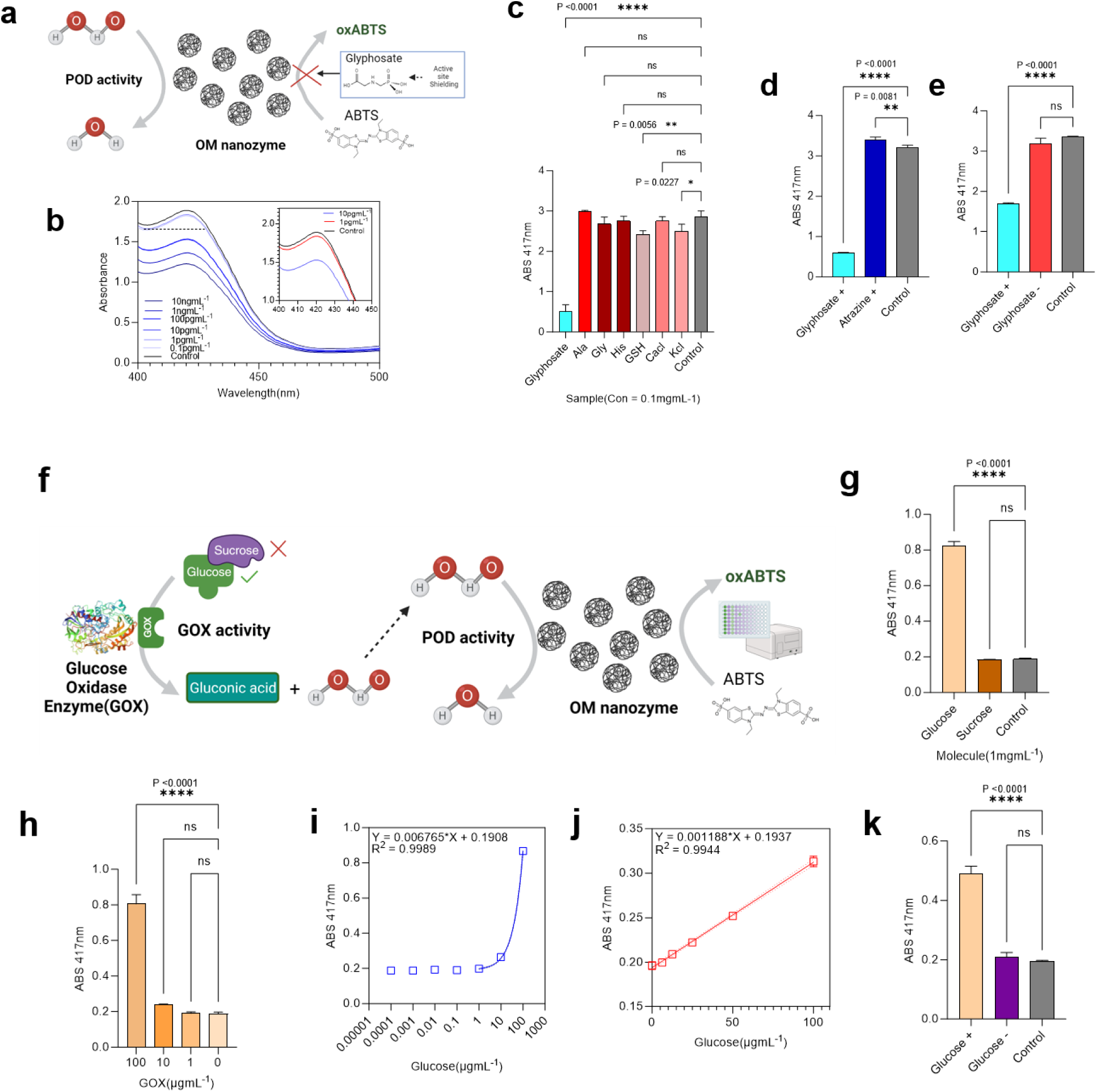
OM nanozyme integrated colorimetric sensing platform for agricultural & biological molecules detection. **a.** Schematic of the OM nanozyme-based glyphosate sensing. **b.** Serial dilution/practical LOD identification of glyphosate(for analytic sensitivity). **c.** Analytic selectivity confirmation(N=3, **** = P<0.0001). **d.** Selectivity confirmation with the comparison with relative herbicide(e.g., atrazine)(N=3, **** = P<0.0001). **e.** Mixture selectivity(N=3, **** = P<0.0001)(related to the analytic specificity). **f.** Schematic of the OM nanozyme-based glucose sensing. **g.** Sugar selectivity, comparison to Sucrose (N=3, **** = P<0.0001). **g.** Gox concentration confirmation (N=3, **** = P<0.0001). **i.** Serial dilution of glucose solution for the LOD identification(for analytic sensitivity) **J.** Concentration-dependent linearity identification for glucose sensing(N=3) **k.** Mixture selectivity (N=3, **** = P<0.0001).

Similar to glyphosate detection, glucose detection performance via OM nanozyme-based colorimetric sensing platform was evaluated (**Figure 4 f-k**). The previously confirmed enzyme-cascade method was applied for glucose detection [7]: In this process, glucose oxidase(Gox) consumes the glucose as the substrate and produces gluconic acid and hydrogen peroxide as a byproduct. The OM nanozyme, in conjunction Gox, consumes hydrogen peroxide as a target substrate and leading to the oxidation of the coloring substrate. Therefore, the absorbance signal generated by the oxidation of coloring substrates is directly correlated to the initial glucose concentration. Optimization of the Gox concentration and sugar selectivity analysis was conducted to establish the optical conditions for the glucose sensing platform (**Figure 4 g, h**). The analytic sensitivity, selectivity, and mixture selectivity analysis were also conducted (**Figure 4 i-k, SI Figure 12**). An affordable LOD of 3.47 ngmL^-1^ (based on the lowest distinguishable concentration value, 3σ,) was achieved along with selectivity and specificity.

### 3.5 OM nanozyme-based, Point-Of-Use biomolecules sensing platform

To narrow the gap between lab and real-world applications in molecule sensing, a PoU platform was developed and integrated with OM nanozyme, highlighting its practicality. Initially, the colored signal was analyzed using UV-VIS spectroscopy; however, an alternative readout system was developed to replace UV-VIS spectroscopy (Figure 5 a). The colorimetric assay’s strength lies in the fact that color intensity indirectly provides analyte concentration, so optical values (e.g., H, S, V) were used instead of absorbance values. Following the colorimetric assay, images were captured using a smartphone camera. A MATLAB-based app was designed to collect optical values for each well plate (**Figure 5 a**). Then, polynomial regression analysis was carried out to create a standard curve for the three-target molecules: H_2_O_2_, glyphosate, and glucose. The linearity of the results was confirmed, yielding reliable profiles using the engineered formulas: I = (0.5*H+0.5*V) for H_2_O_2_, I= (0.8*H+0.2*V) for glyphosate, and I= (0.7*S+0.3*V) for glucose (**Figure 5 b-d**), respectively. To prove its potential as an alternative analytic tool to conventional analytic tools (e.g., LC-MS/HPLC), sample measurements were partially conducted. In LC-MS analyses, a non-derivatization method was used to directly compare with the colorimetric assay, as this method does not require any pre-prep steps for the sample measurement. While the analytic limit of LC-MS decreased due to the quality of non-derivatization samples, the same sample conditions were applied to the colorimetric assay to ensure a genuine analytical comparison (**SI Figure 13**). In HPLC sample analysis, syringe filtering is necessary to prevent clogging, which can lead to a decrease in target molecule concentration. Considering these factors, the OM-nanozyme-based biomolecule sensing platform demonstrated superior analytical sensitivity and practicality compared to these traditional tools (**Figure 5 e, SI Figure 14**). Furthermore, μPADS were designed to finalize the PoU platform (**Figure 5 f**). The μPADS dimension was confirmed as around 30mm × 30 mm, using chromatography-grade cellulose paper. A simple, round-shaped well design was selected, mimicking the wells in a standard well plate, with a capacity to hold up to 15uL of solution. Multiple replicable wells were located around the main spot (deduced dimension) and a hydrophobic polymer coating was conducted around each well to promote phase difference. (**Figure 5 g**). The Z-height gap between the wells and the borders was measured using confocal microscopy and Raman microscopy, showing a difference of more than 20 micrometers (Z-axis)(**SI Figure 15,16**). SEM analysis was conducted to examine the μPADS surface, and each part of μPADS was observed visually. To confirm the presence of the OM nanozyme residue on the μPADS, an SEM image was obtained (**SI Figure 17**), followed by EDS mapping to verify the existence of the OM nanozyme (**Figure 5 h**). Although the smartphone-based image readout and related regression analysis system have lower analytic sensitivity compared to the conventional UV-VIS spectroscopical readout, it is still at a usable level (**SI Figure 18**). Thus, this procedure was applied to the μPADS system’s optical value analysis. The smartphone image was obtained similarly to the previous steps (**Figure 5 i**) and regression analysis was conducted to fit the data. A new fitting model and formula were established showing satisfactory R2 (R^2^> around 0.8 or more) for H_2_O_2_, glyphosate, and glucose concentrations (**Figure 5 j,k, SI Figure 19-22**). The practical LOD was measured as at least 1.6μgmL^-1^ for glyphosate and for glucose, respectively. Therefore, the OM enzyme-based PoU sensing platform demonstrated its potential for agricultural and biological molecule detection in a lab-free environment.

**Figure 5.**
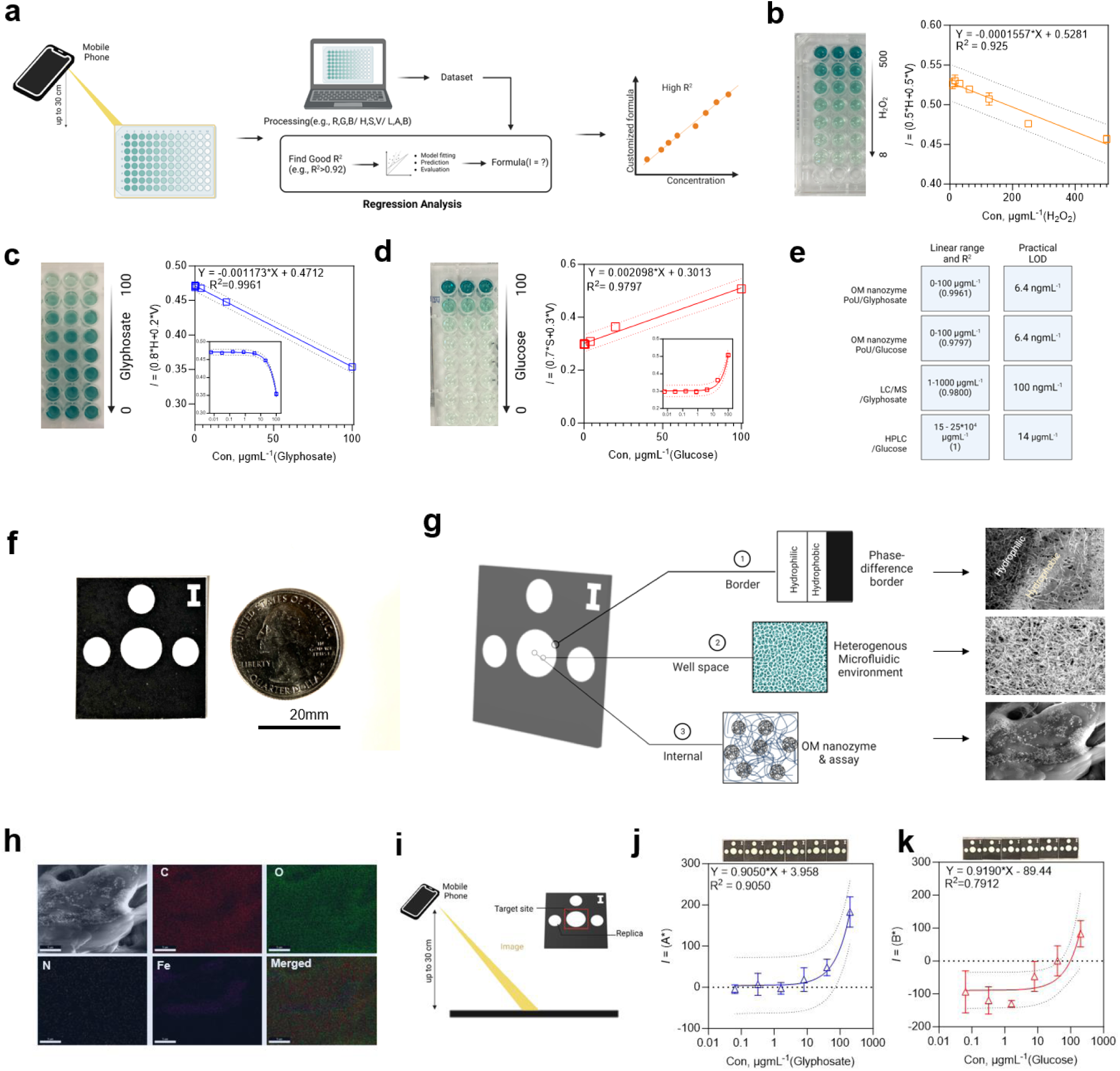
OM nanozyme-integrated, Point-of-Use application platform for dual molecule detection. **a.** Schematic of the smartphone-based sensing procedure incorporated fitting process (the regression analysis, polynomial). **b.** Concentration-dependent linearity profile by regression-analysis methods (molecule: H_2_O_2_, image obtained by smartphone, N=3). **c.** Concentration-dependent linearity profile by regression methods (molecule: glyphosate, image obtained by smartphone, N=3). **d.** Concentration-dependent linearity profile by regression methods(molecule: glucose, image obtained by smartphone, N=3) **e.** compassion of the molecule sensing performance with UV-VIS spectroscopical methods(with OM nanozyme) and conventional analytic tools(e.g., LC/MS, HPLC, etc.).**f.** Real image of a μPADS for the assay(scale, 20mm) **g.** Schematic illustration of the μPADS with the location, and the corresponding SEM image. **h.** EDS mapping image of the well part (sensing part) of the μPADS (OM nanozyme-located) **i.** Graphical illustration of the smartphone-based molecule sensing procedure on a μPADS. **j.** linearity analysis is driven by regression analysis for glyphosate Top: the real image of the assay on the chip. **k.** linearity analysis is driven by regression analysis for glucose Top: The real image of the assay on the μPADS.

## 4. Conclusion

In this study, a novel organic compound and monomer-based, sustainable nanozyme (OM nanozyme) was introduced that exhibits a decent peroxidase-like catalytic activity. The advanced, double-chelation-based fabrication methodology provides enhanced feasibility of nanozyme in terms of its smaller physical dimensions with sustainable characteristics including degradability for partially resolving further waste management concerns. The OM nanozyme exhibits decent enzyme-like catalytic performance (peroxidase-like activity), as *Km* = 0.046mM (H_2_O_2_). The OM nanozyme-based customized molecule sensing system enables the detection of dual agricultural (e.g., glyphosate) and biological molecules (e.g., glucose), with decent analytic sensitivity. It also demonstrates the capability to detect pseudo-real samples prepared from agricultural/biological molecule mixtures. The integration of a smartphone-image based readout and a paper microfluidic chip (μPADS)-based hardware with OM nanozyme highlights its potential for on-demand molecule sensing in lab-free environments. This sustainable OM nanozyme with its PoU sensing system is envisioned for direct applications in agriculture, food, and biologically in the future.

## Supporting information

Supporting list

## Supporting Information

The supporting information can be found below[from page 27].

## Acknowledgements

We appreciate the experimental and research support provided by the staff of ITG (Imaging Technology Group), Beckman Institute, Material Research Laboratory, High Throughput Screening Facility (utilization of BioTek Cytation 5 Multi-mode Imaging Reader, funded by the office of the Director, National Institutes of Health, award #S10 OD025289), EPR laboratory, Mass Spectrometry Lab and Integrated Bioprocessing Research Laboratory at the University of Illinois at Urbana-Champaign.

## Abbreviations

OM nanozyme: Organic compound/monomer based, sustainable nanozyme
Nanozyme: Nanomaterial-based enzyme mimetics
POD activity: Peroxidase-like activity EPR, Electron paramagnetic resonance
ABTS: 2,2’-azino-bis (3-ethylbenzothiazoline-6-sulfonic acid
TMB: 3,3,5,5’-Tetramethylbenzidine
OPD: o-phenylenediamine
SEM: Scanning electron microscopy
TEM: Transmission electron microscopy
FTIR spectroscopy: Fourier transform infrared spectroscopy
XRD: X-ray Diffraction
XPS: X-ray photoelectron spectroscopy
LC-MS: Liquid chromatography-mass spectrometry
HPLC: High-performance liquid chromatography.

## Supporting Figures

**SI Figure 1.**
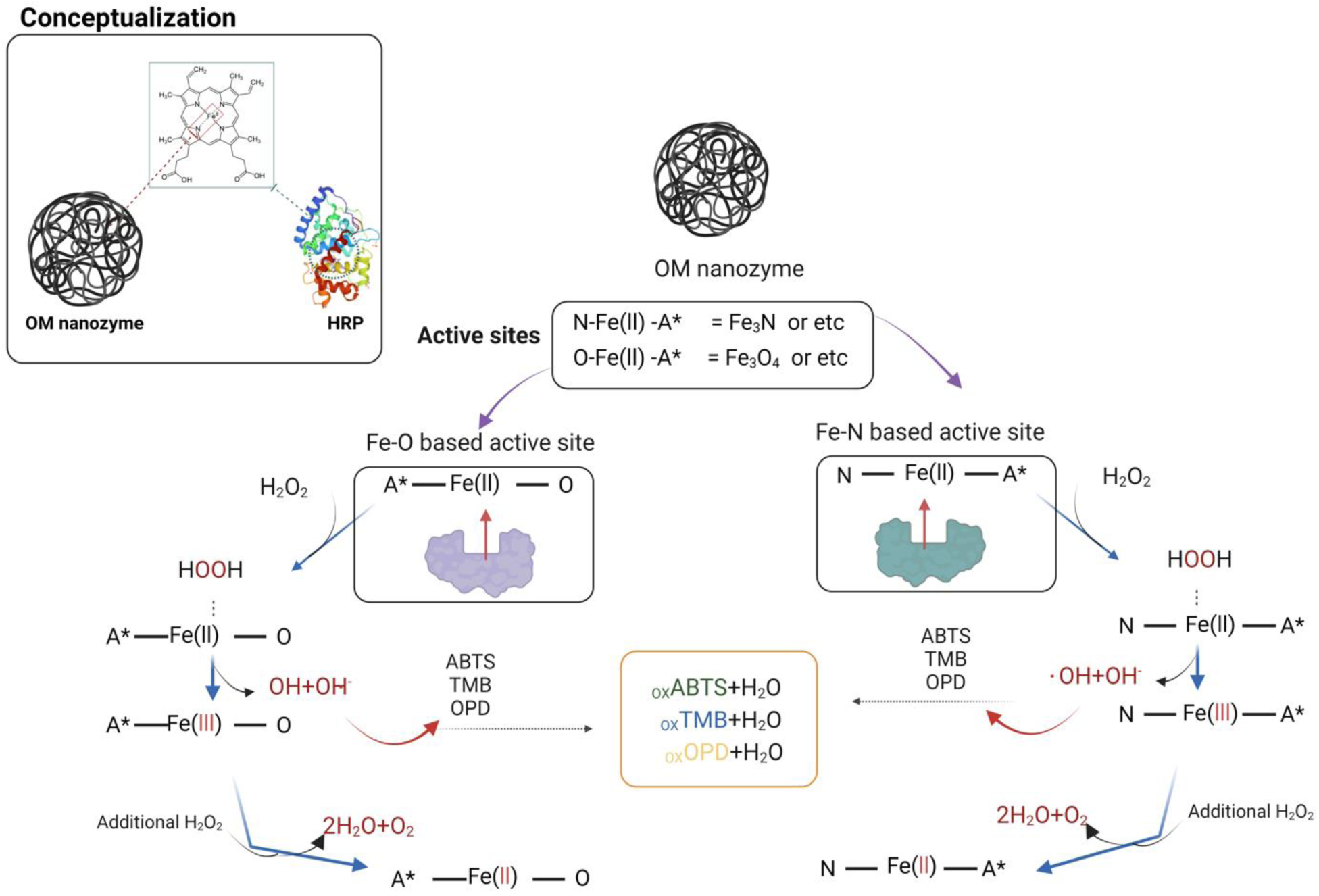
Graphical illustration of the conceptualization of OM nanozyme and its expected enzyme-like catalytic pathway. Fe_x_-O_y_, and Fe_x_-Ny (x and y <6) sites are OM nanozyme’s designated active sites.

**SI Figure 2.**
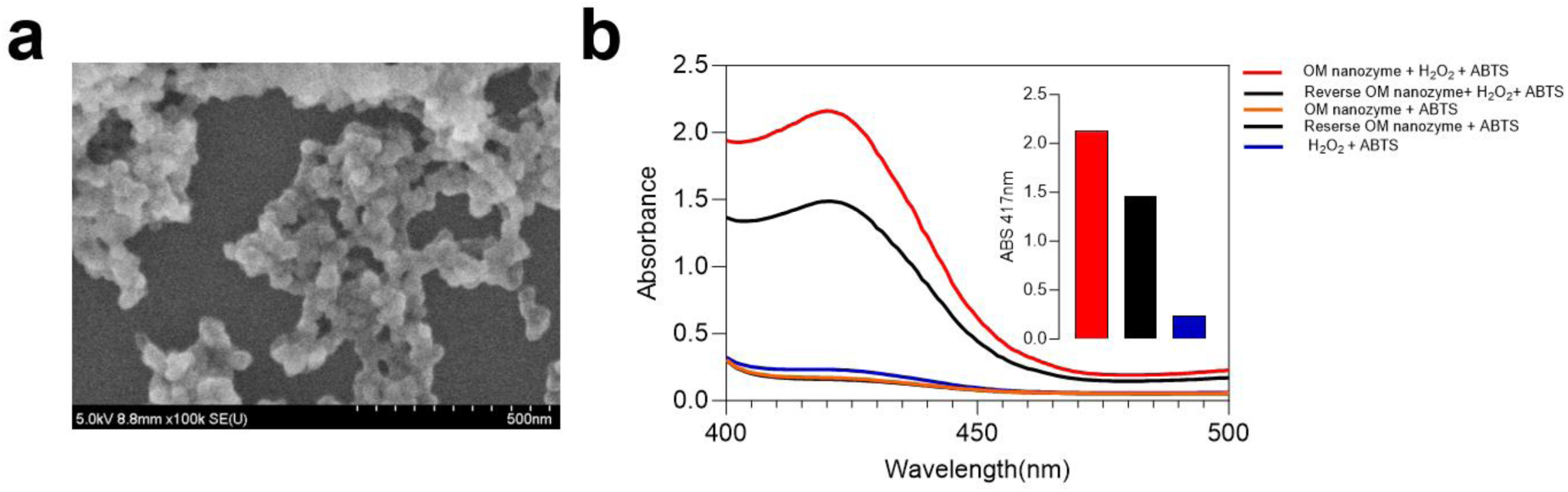
Confirmation of the fabrication step configuration. a. SEM image of reversed fabrication/configuration (fabrication starting from monomer, not organic compounds) b. Absorbance peak comparison between OM and the reversed OM for the confirmation(Inserted bar graph, ABS peak at 417nm, OM > Reversed OM).

**SI Figure 3.**
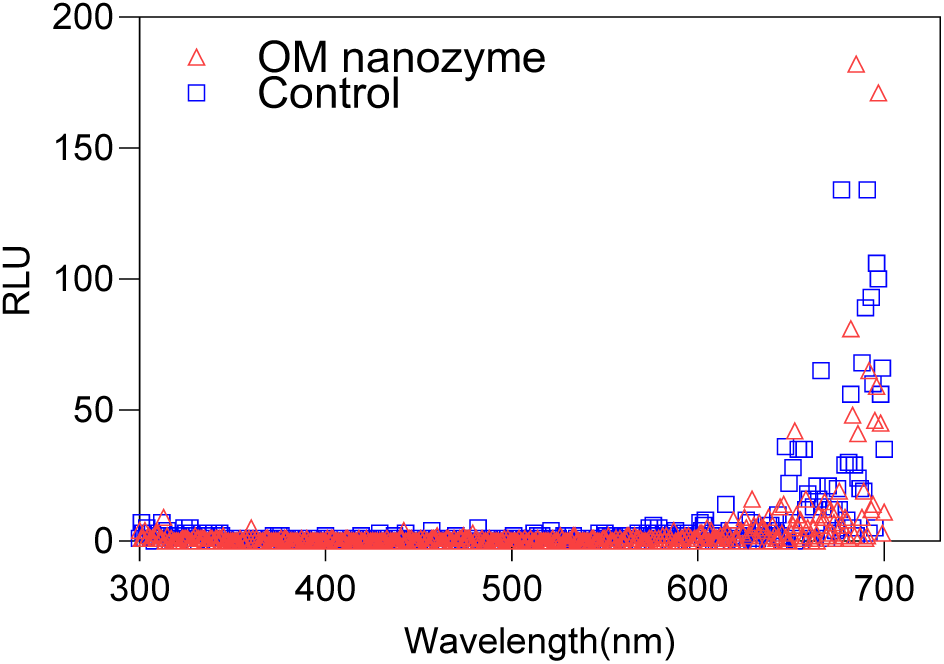
Luminescence spectroscopical scanning profile of OM nanozyme Red: OM nanozyme Blue: control(buffer).

**SI Figure 4.**
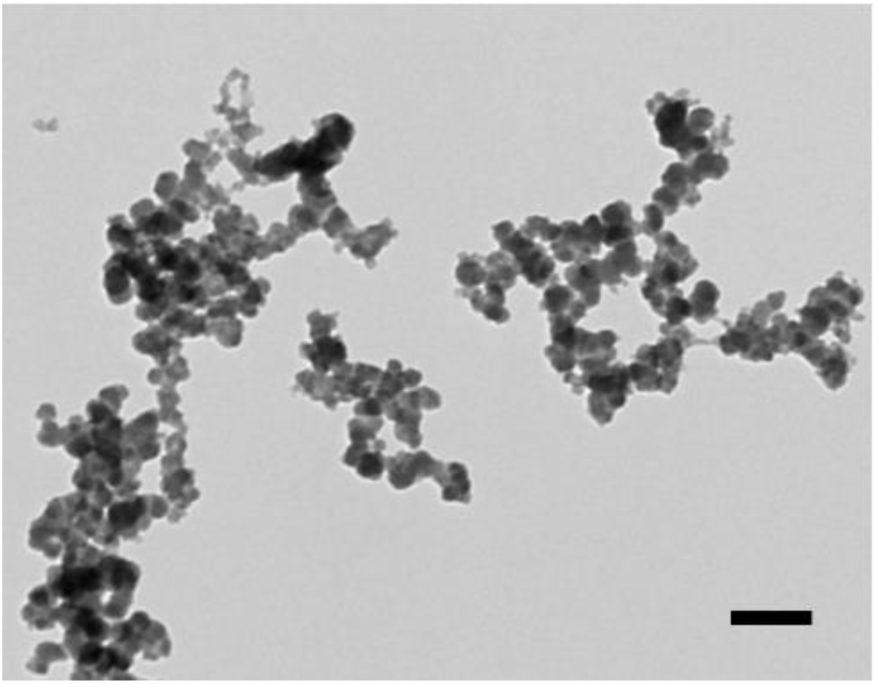
Additional TEM image of OM nanozyme (scale = 100nm).

**SI Figure 5.**
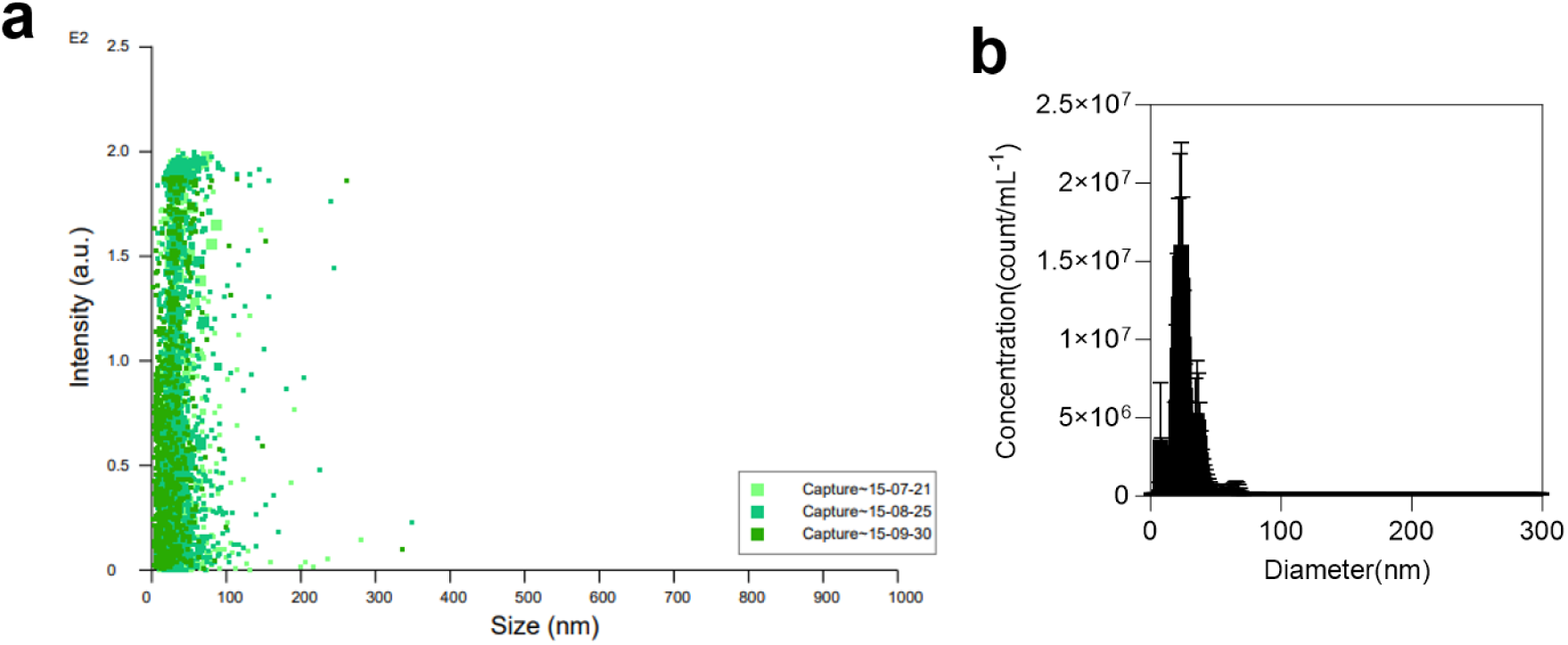
NTA (nanoparticle tracking analysis) of OM nanozyme. a. Raw profile of the particle distribution, 3 cycle. b. translated size distribution graph.

**SI Figure 6.**
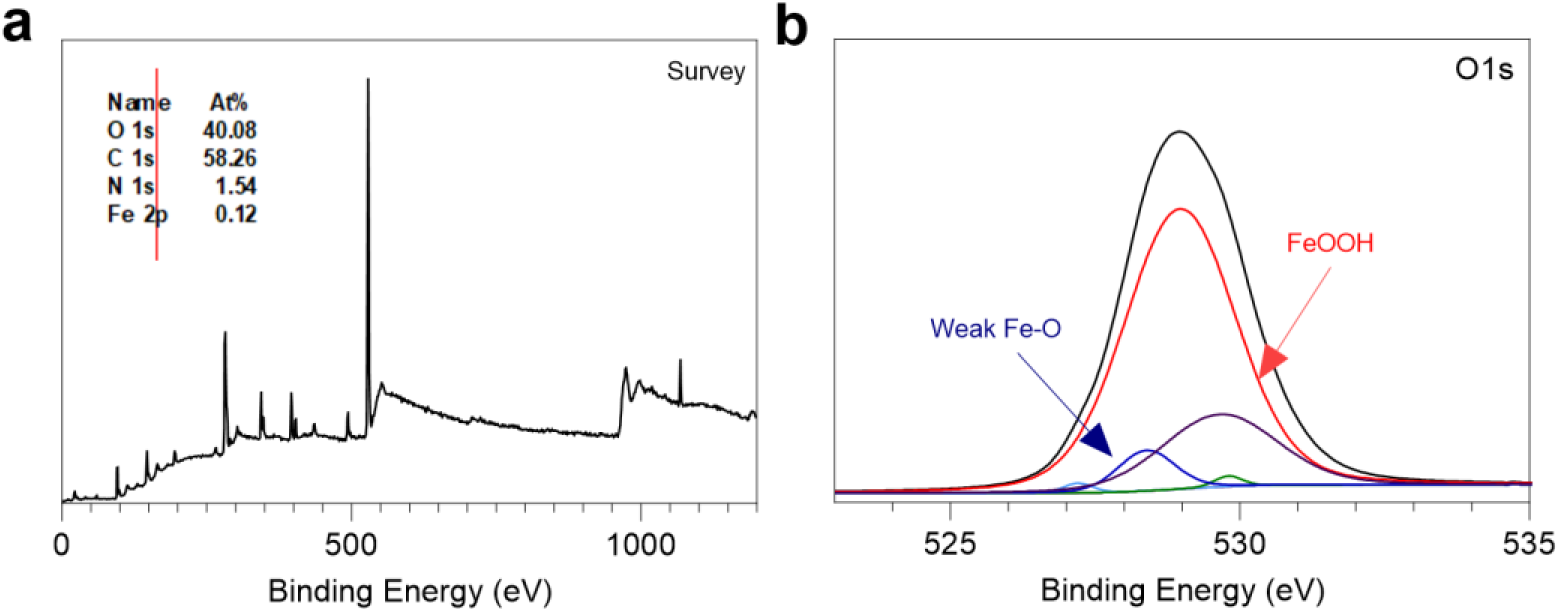
XPS quantification and deconvolution information at the O 1s(range, 523 eV to 535 eV). Purple : 528.39eV(weak Fe-O bonding)[SI ref 7], Red: 528.96-97eV(Fe-O((FeOOH))[SI ref 8], Green: 529.79eV(O^2-^)[SI ref 9].

**SI Figure 7.**
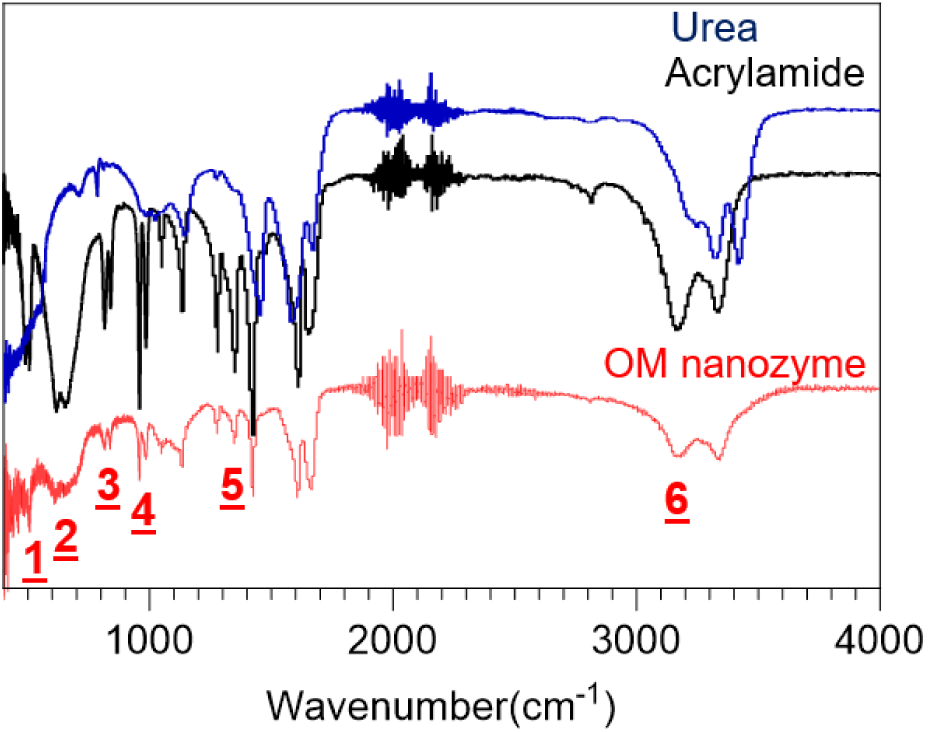
FT-IR analysis of the OM nanozyme. Red: OM nanozyme Blue: Urea Black: Acrylamide (Around 2000 cm^-1^, instrumental noise). [1]. 438 cm-1 (δC = C–O, from Urea) [SI Ref 1] [2]. 575 cm-1 (Fe-O)) [SI Ref 2] [3]. 719.1 cm(NH band)) [SI Ref 3] [4]. 882 cm-1 (Fe-N)) [SI Ref 4] [5]. 1431 cm-1 (Bending(deformation) vibration of C-H)) [SI Ref 5] [6]. 3210 cm-1 (N-H stretching)) [SI Ref 6]

**SI Figure 8.**
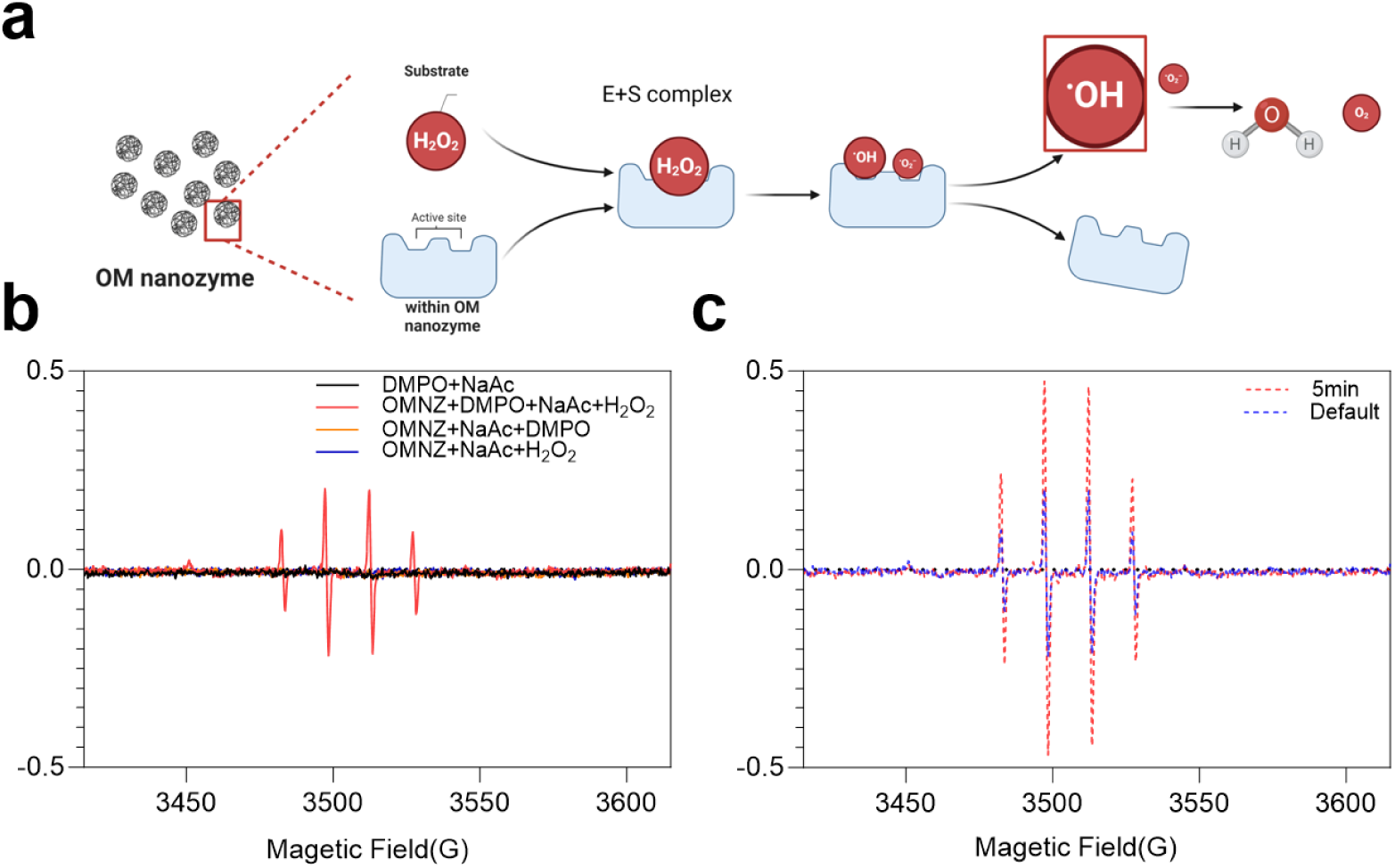
EPR analysis of OM nanozyme. a. Expected Enzyme-like catalytic mechanism of OM nanozyme and byproducts(e.g., hydroxyl radical, before being H_2_O and O_2_). b. overlapped c. indirect kinetics via EPR analysis, Red: 5 minutes after measurement Blue: Default.

**SI Figure 9.**
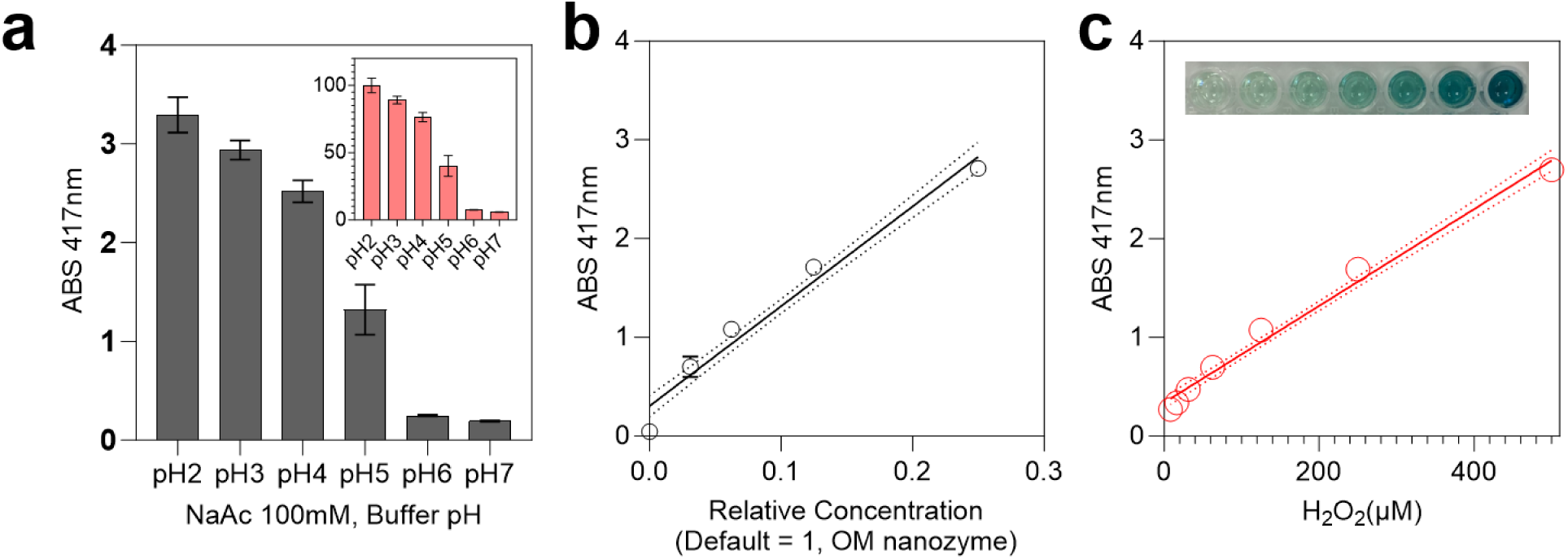
Optimization and characterization of experimental condition of substrate-dependent colorimetric assay. a. pH dependent(N=3), b. OM nanozyme’s concentration dependent(Y = 10.11X + 0.3026, R^2^=0.9654, N=3) and c. the substrate-concentration response correlation of absorption signal(Y = 0.004902X + 0.3400, R^2^=0.9868, N=3).

**SI Figure 10.**
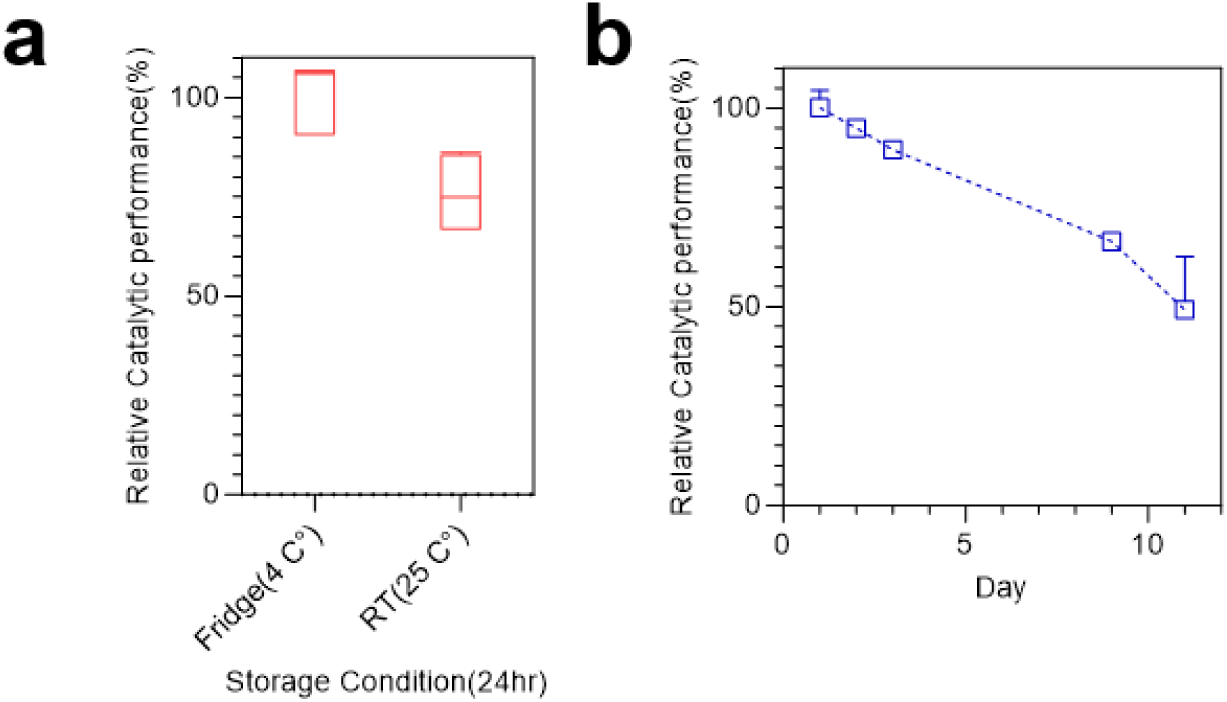
Storage stability of OM nanozyme a: temperature dependent comparison b: relative enzyme-like catalytic activity validation through storage period. (Half-life, 11days/approx., N=5).

**SI Figure 11.**
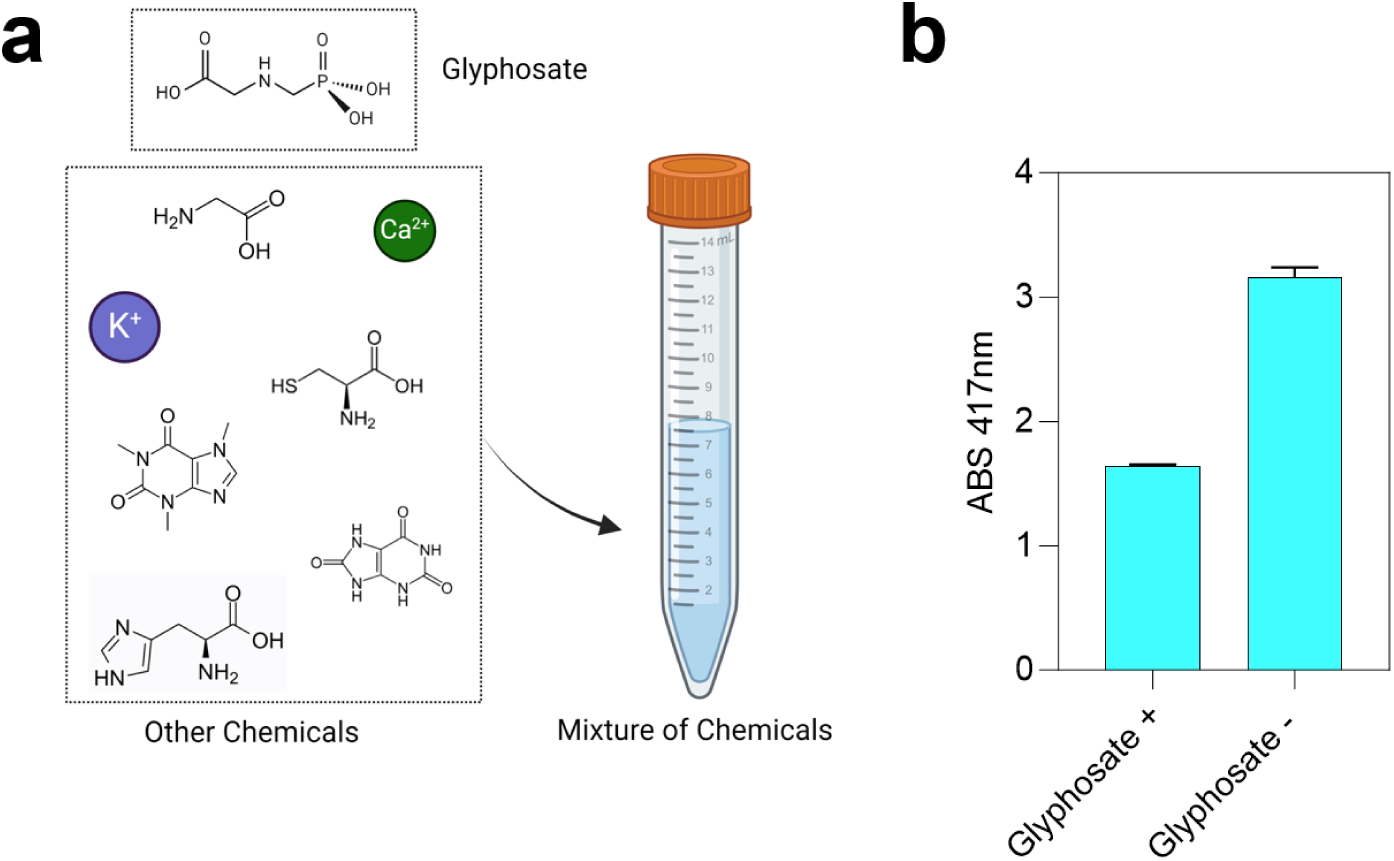
Schematic of the validation of the mixture selectivity of glyphosate. (Concentration, at least 1mgmL^-1^ N=3) a. Graphical illustration of the mixture solution(with 6+ agricultural/biological molecules) b. Absorbance measurement of two samples for the comparison.

**SI Figure 12.**
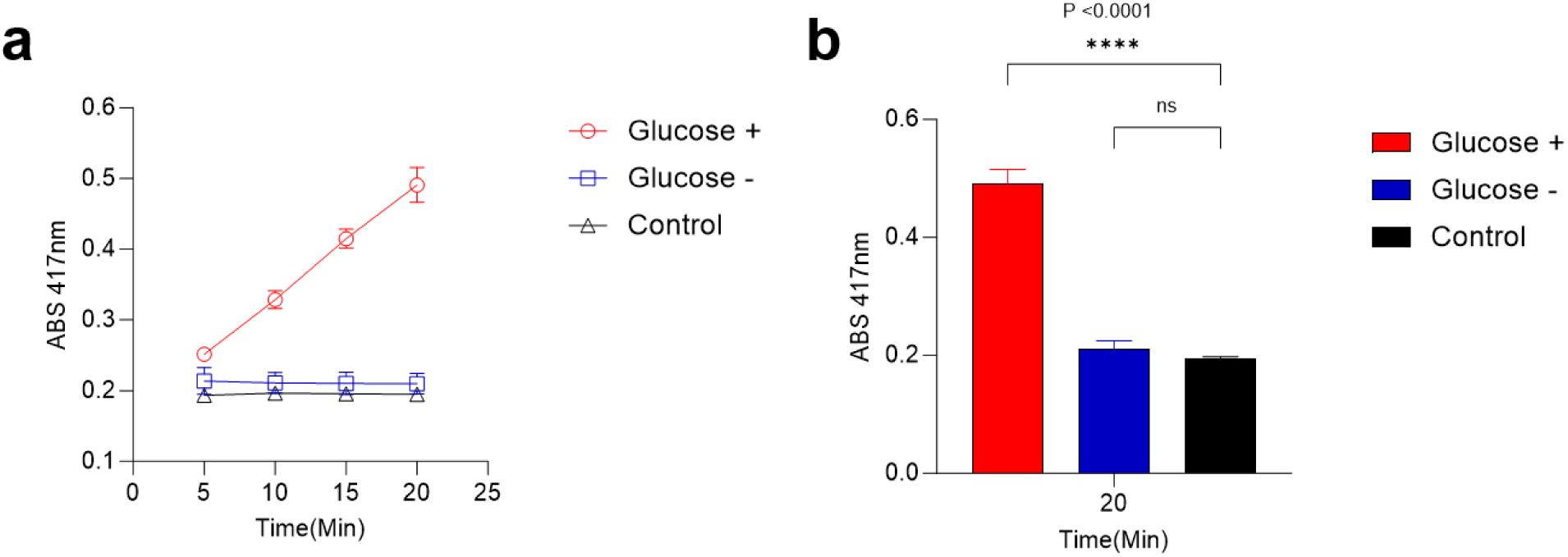
Time-dependent absorption compassion of the mixture selectivity of glucose (time period, 5 minutes to 20 minutes, 5-minute term, **** = P<0.0001, N=3) a. Time-dependent absorption comparison(N=3), b. Statistical analysis (one-way ANOVA) of confirmation related to the selective glucose detection on the mixture solution.

**SI Figure 13.**
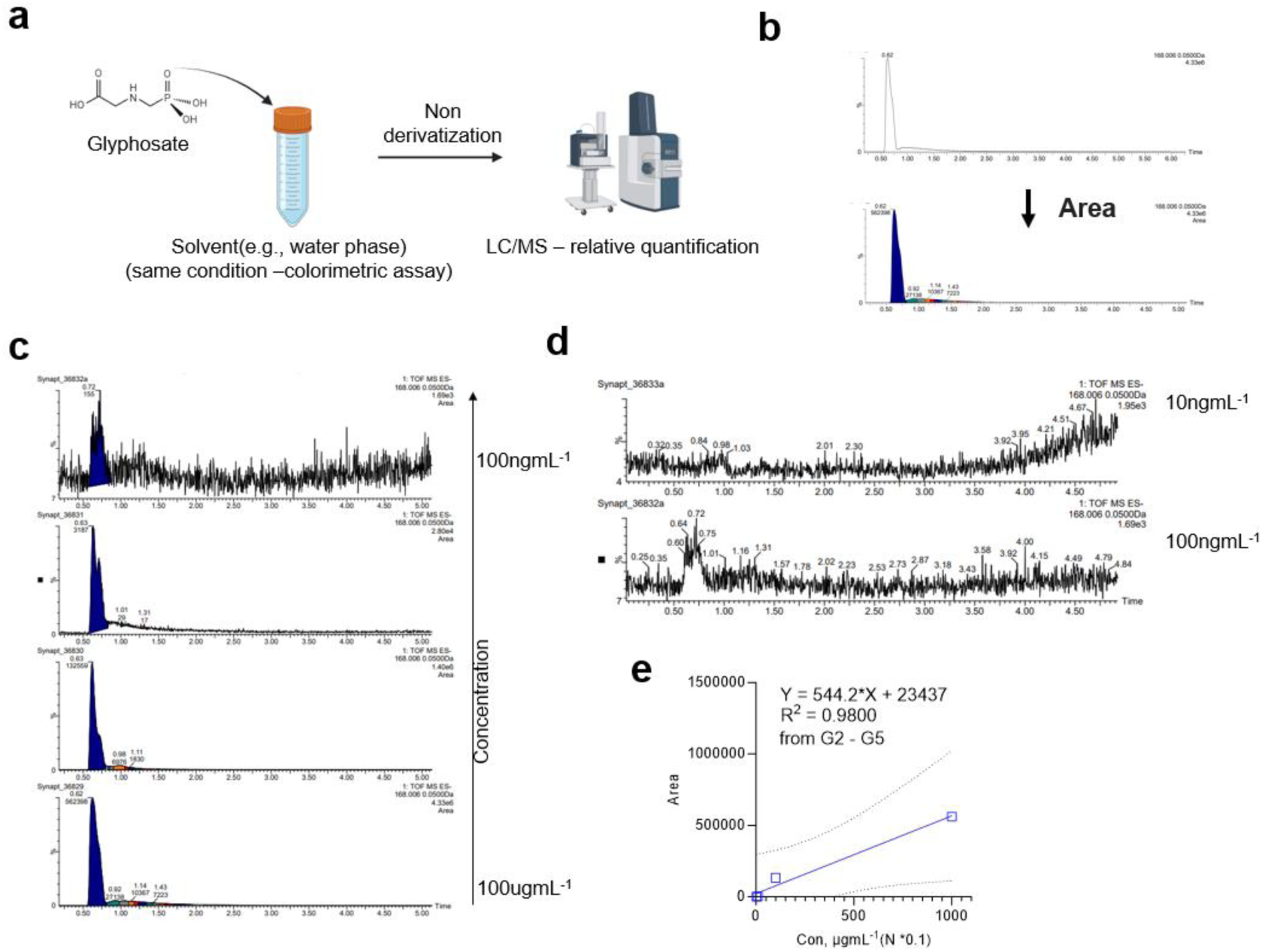
Compassion of the glyphosate detection system with the conventional analytic tool (e.g., LC/MS with non-derivatization methods/crude mode) a. Schematic of the experimental procedure b. examples of quantitative analysis (e.g., area targeting) c. Area measurement depend on the serial dilute glyphosate sample (from 100ugmL^-1^ to 100ngmL^-1^) e. estimated LOD range confirmation (compared to 10ngmL-1) e. translated graph.

**SI Figure 14.**
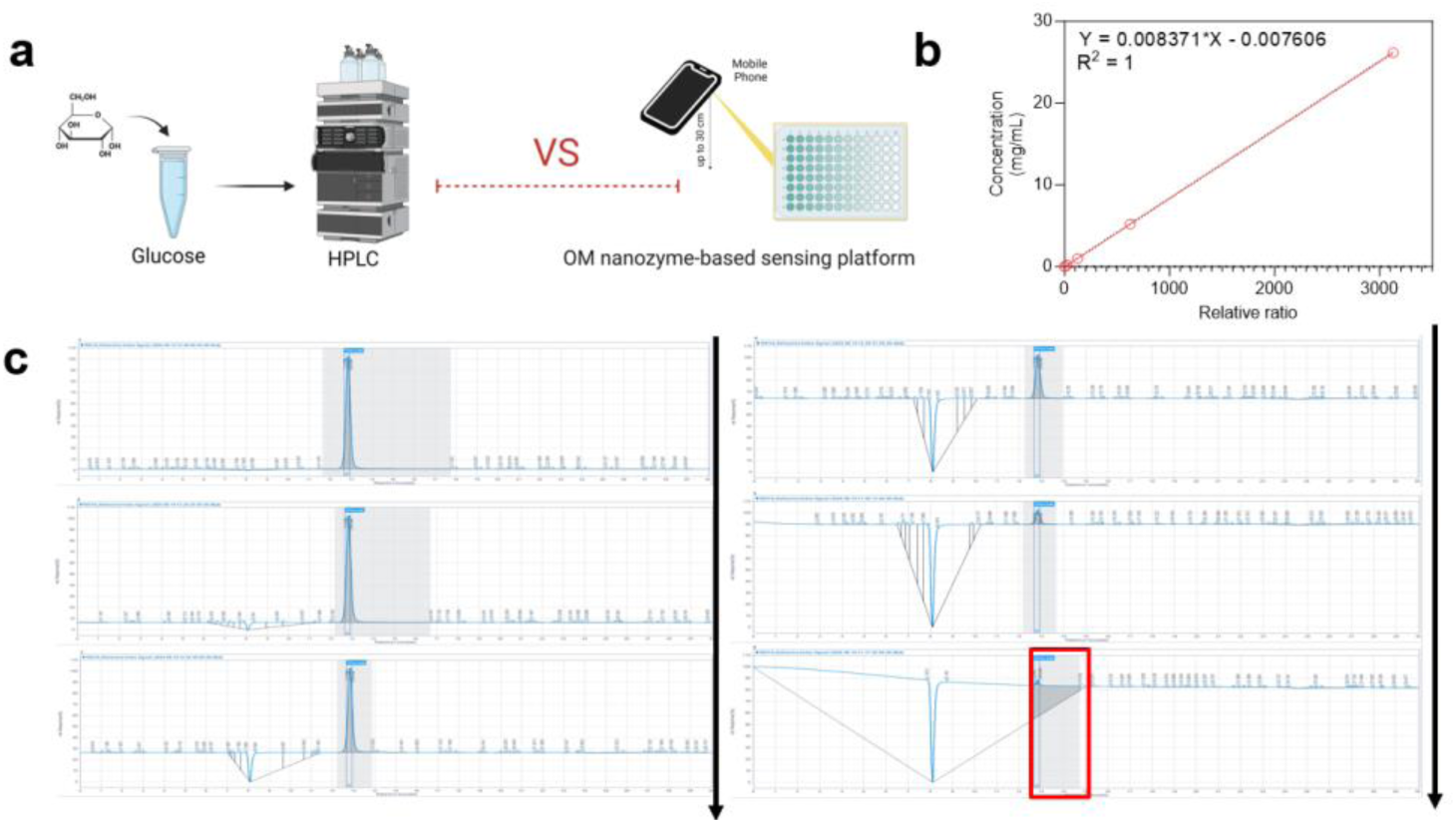
Compassion of the glucose detection system with the conventional analytic tool (e.g., using HPLC) a. Schematic of the experimental procedure b. linearity plot c. Screenshot of the profiles, practical LOD was estimated at least 14ugmL^-1^.

**SI Figure 15.**
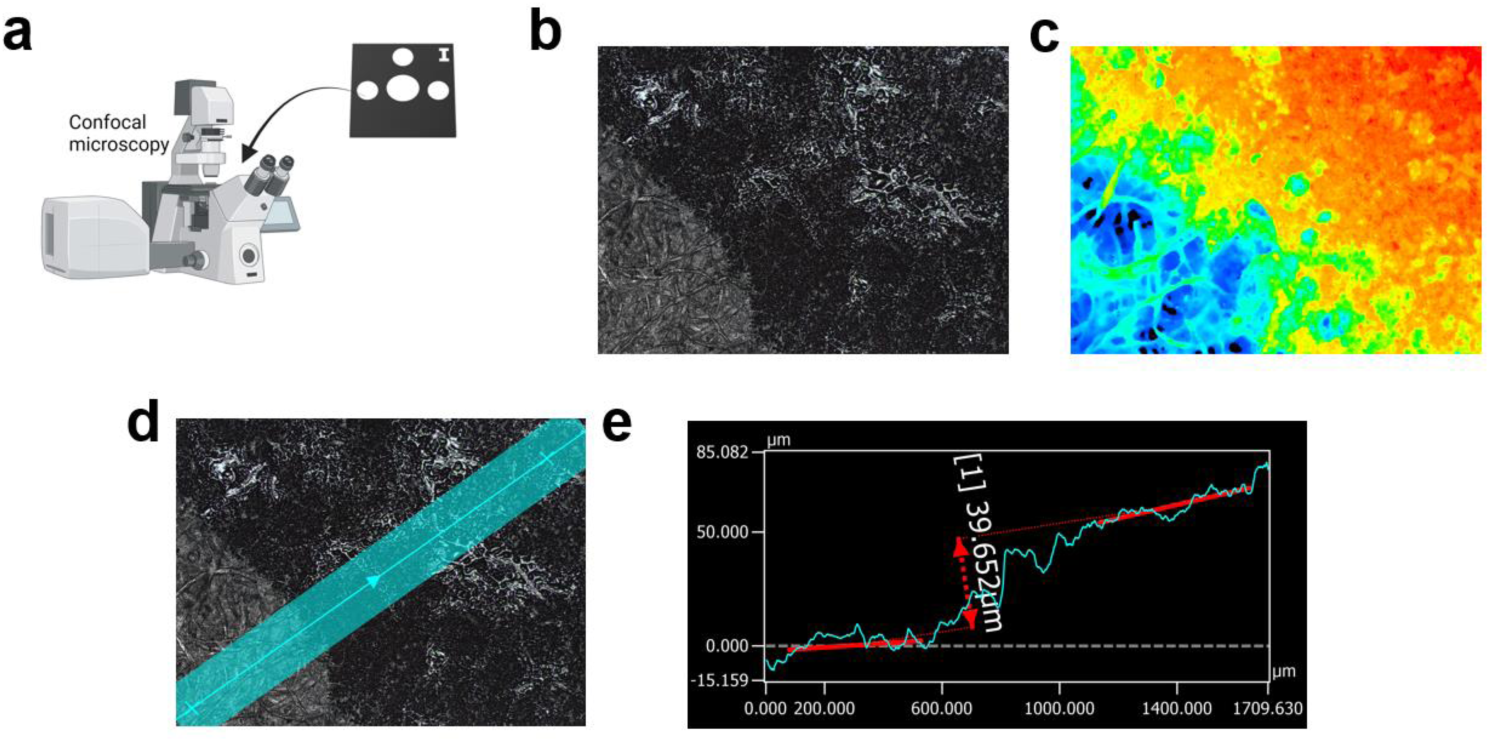
Characterization of the paper-microfluidic chip using confocal microscopy a. schematic of the procedure. b-d. focused image of the well, border and the background, and their measurement line confirmation e: Z-axis analysis provide the gap differences between border and the well.

**SI Figure 16.**
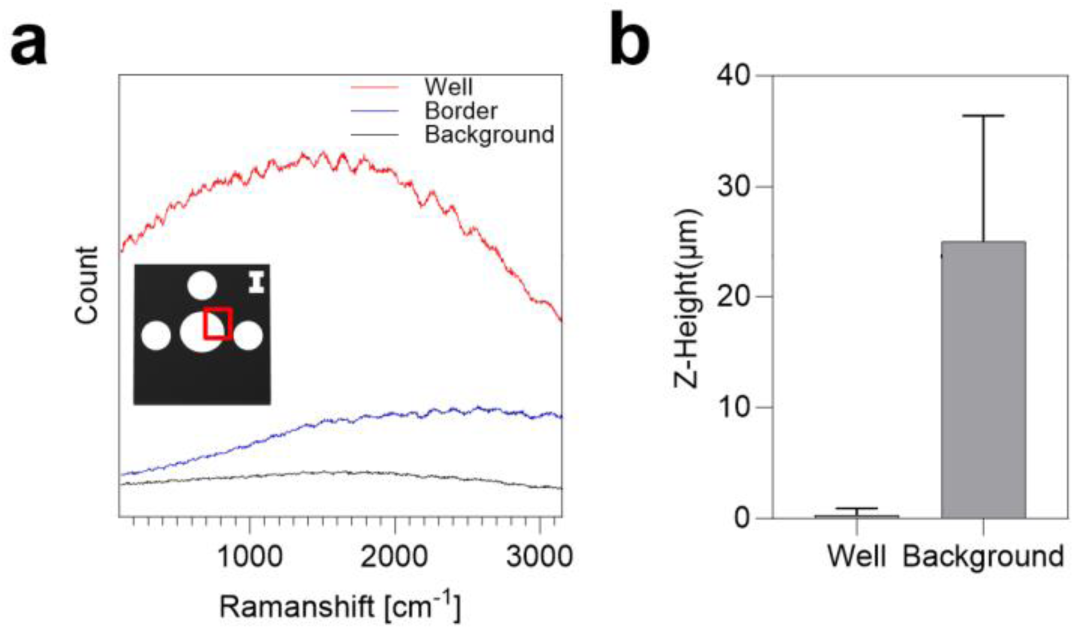
Additional characterization of paper microfluidic chip. a. Raman spectroscopical analysis b. Z-height analysis obtained from confocal microscopy(N=3).

**SI Figure 17.**
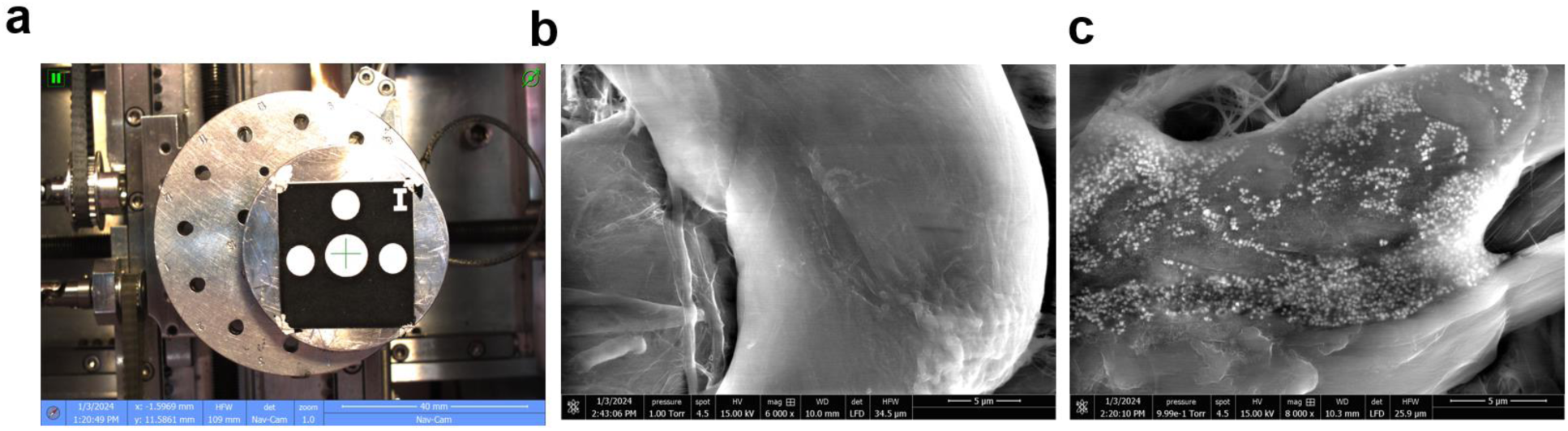
SEM analysis of OM nanozyme mounted on the paper-microfluidic chip. a. target spot b. SEM image of the well spot (control) c. SEM image which OM nanozyme was mounted on the chip in the well spot.(B, C scale = 1μm).

**SI Figure 18.**
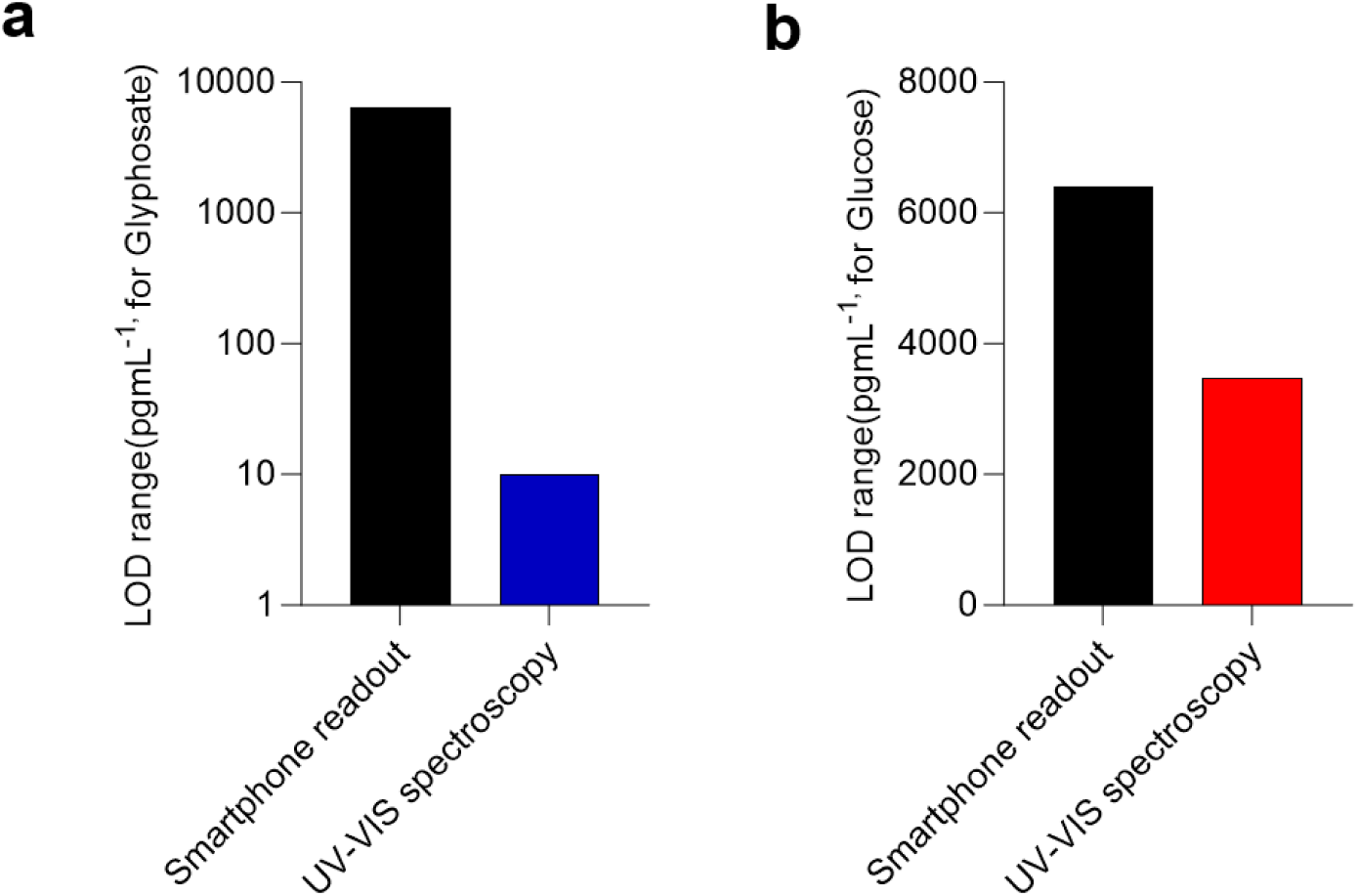
Performance comparison between smartphone readout and the UV-VIS spectroscopy as a readout (Detection limit) a. Glyphosate b. Glucose.

**SI Figure 19.**
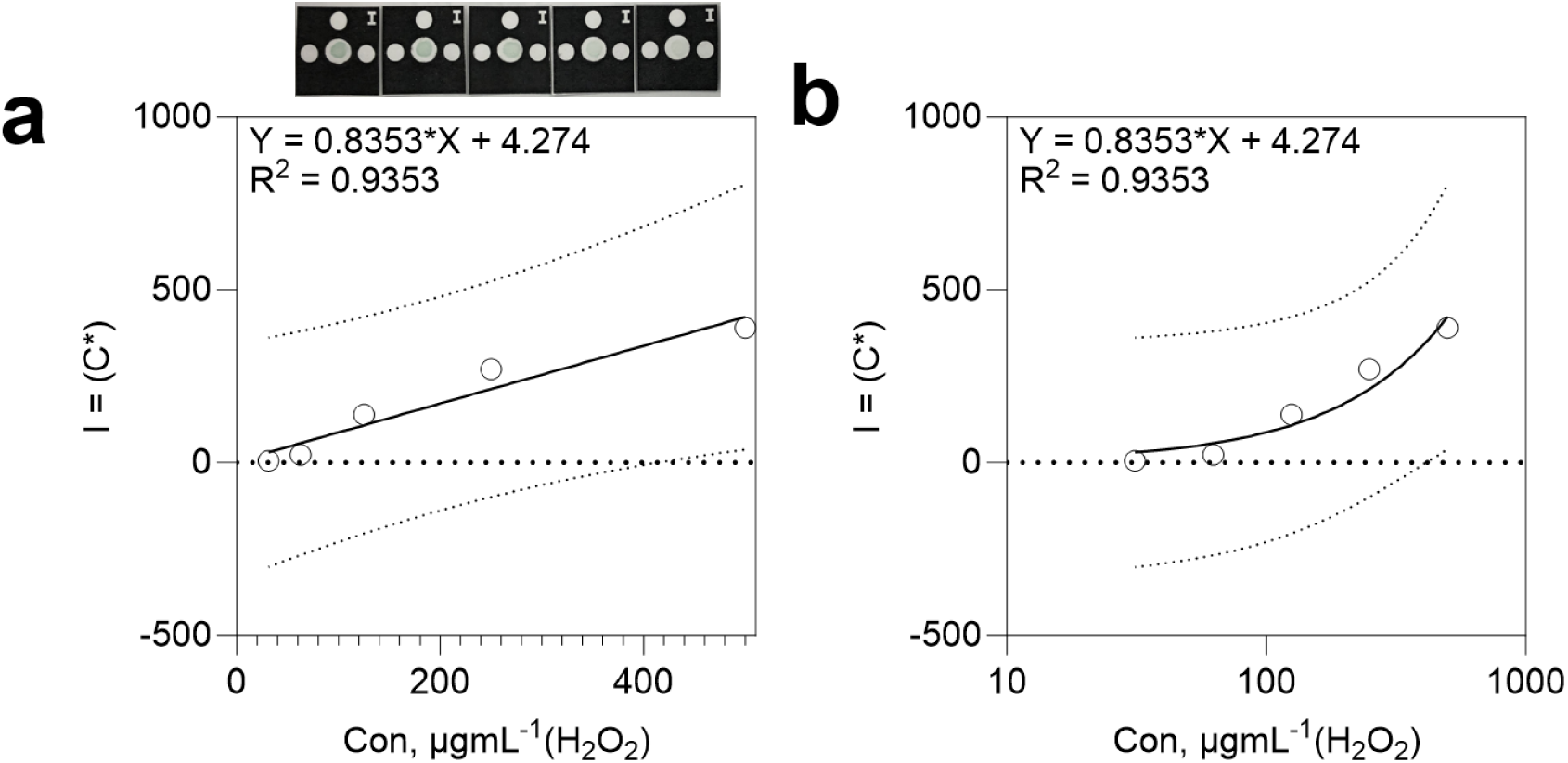
Linear graph of the H_2_O_2_ colorimetric assay/smartphone readout analysis on paper microfluidic chip (above: real image) a. Linear b. Log10 scale (N=3).

**SI Figure 20.**
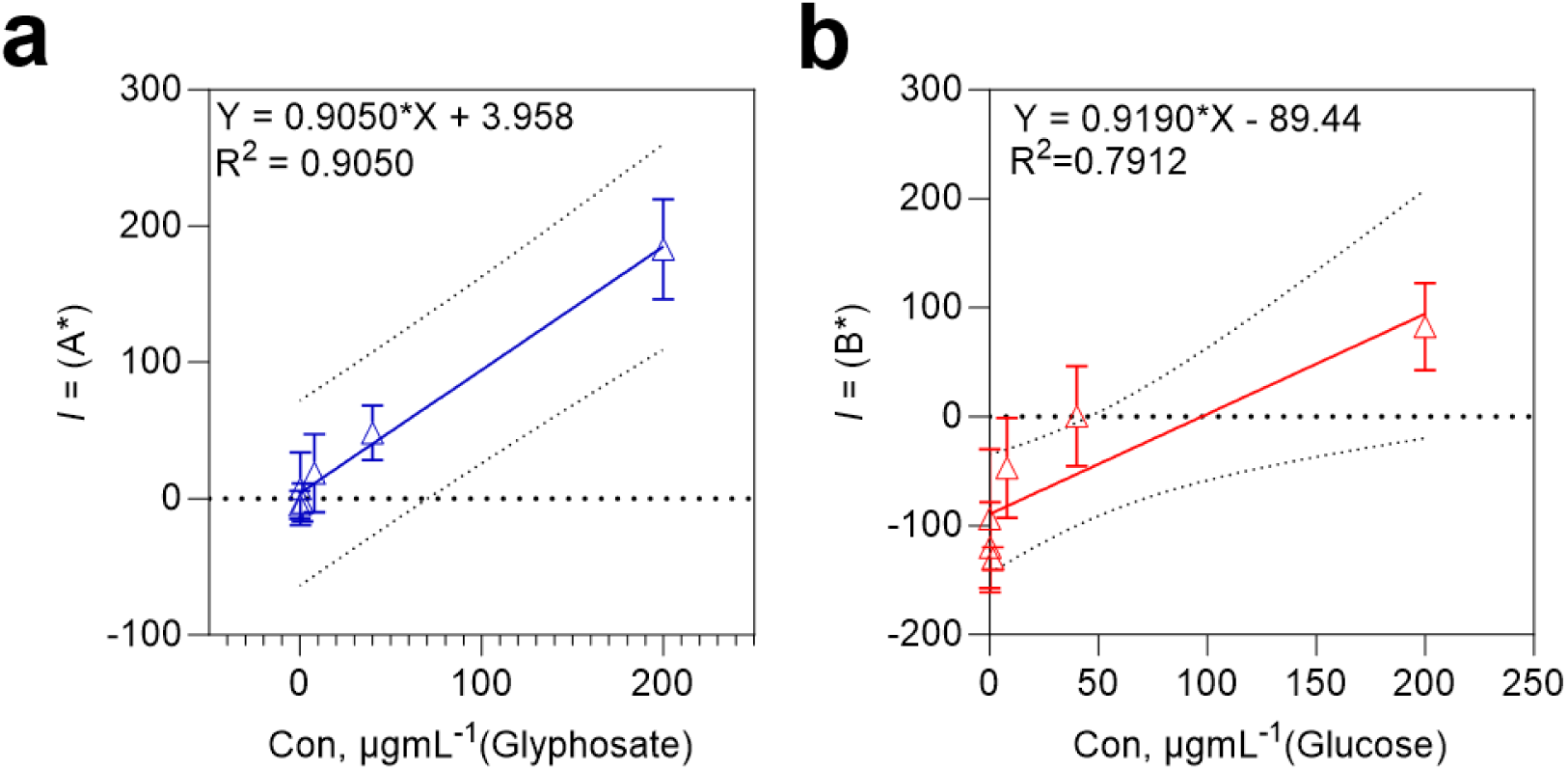
Linear graph of the molecule detection/smartphone readout analysis on paper microfluidic chip a. Glyphosate b. Glucose (N=3).

**SI Figure 21.**
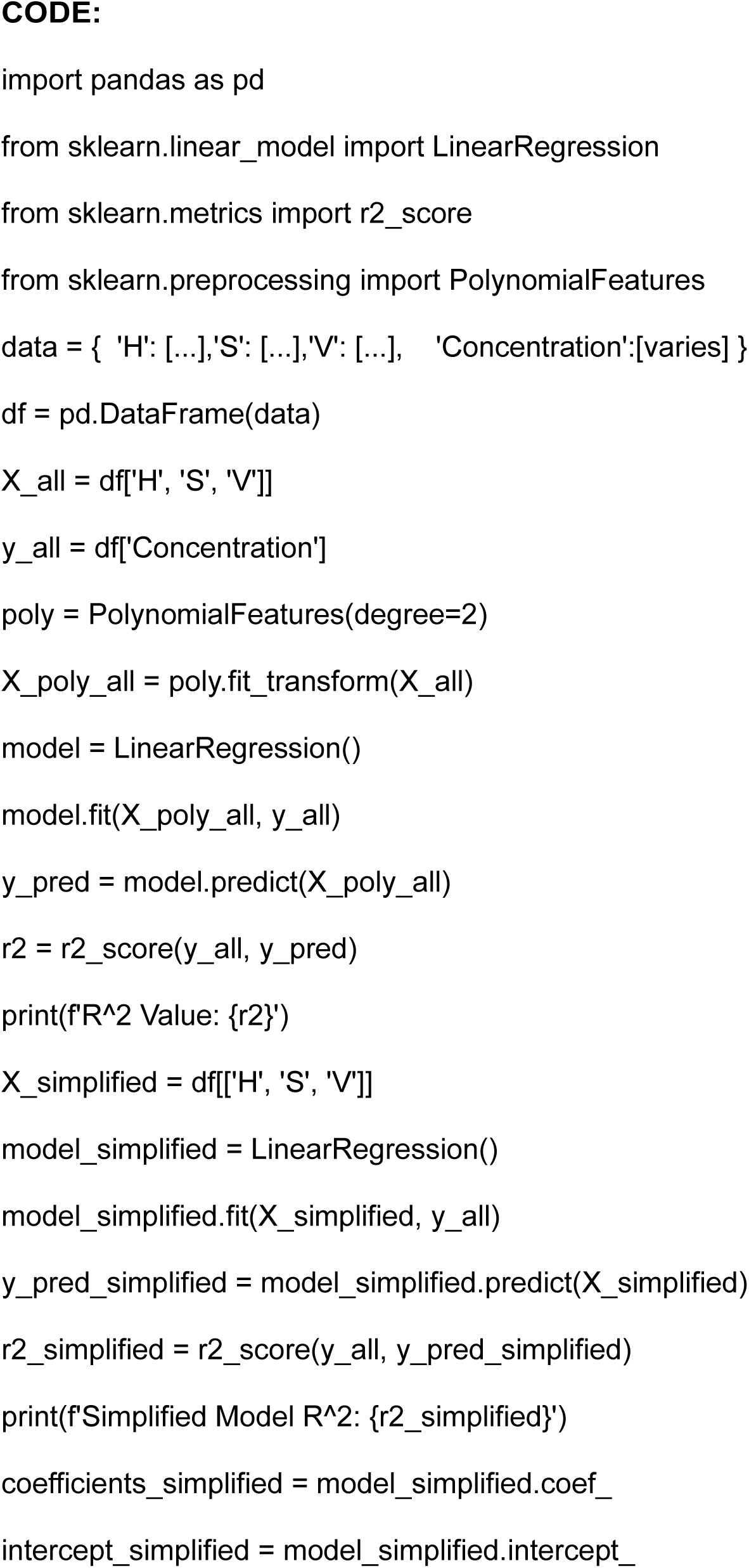
Simplified(partially revised) Python code for obtaining the best R^2^.

**SI Figure 22.**
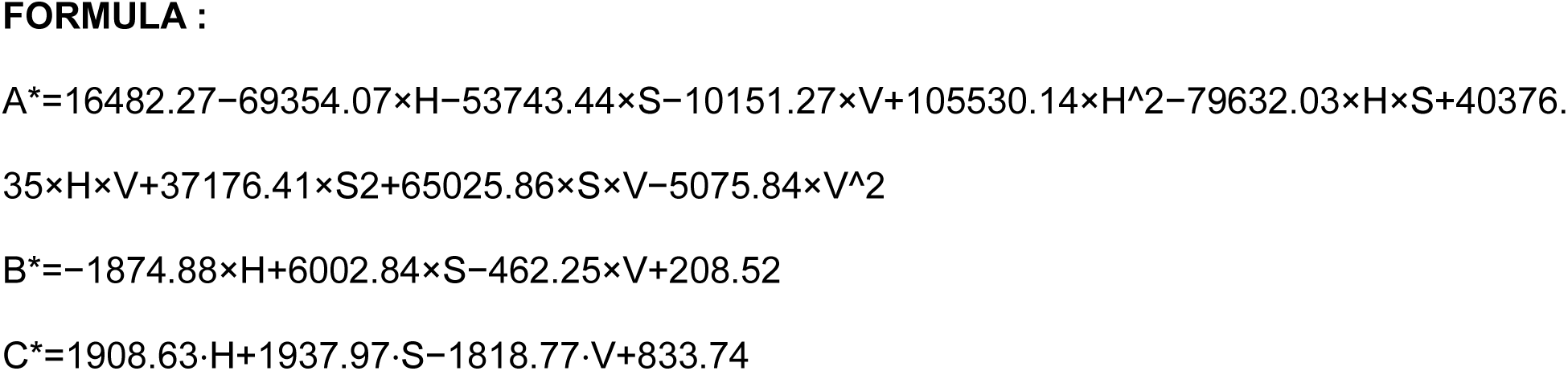
The formula for the fitting (A* = glyphosate, B* = Glucose, C* = H_2_O_2_).

**Table S1.** Kinetic profile table (with other nanozymes)

| Nanozyme | Km(mM) | Vmax<br>($\mu\text{M s}^{-1}$) | Kcat<br>($\text{min}^{-1}$) | Substrate | Reference |
| --- | --- | --- | --- | --- | --- |
| <u>OM</u><br><u>nanozyme</u> | <u>0.046</u> | <u>1.5220</u> | <u>-</u> | <u>$H_2O_2$</u> | <u>This work</u> |
| Co@Fe MOG | 0.72 | 0.028 | - | $H_2O_2$ | Ref.SI.10 |
| Pb-CDs-Fe | 24 | 11.1 | 204 | $H_2O_2$ | Ref.SI.11 |
| Au/Fe-Ag<sub>2</sub>O<sub>2</sub> | 1,16 | 0.871 | - | $H_2O_2$ | Ref.SI.12 |
| Fe<sub>SA</sub>@FibBLG | 0.04 | 0.788 | 21.9 | $H_2O_2$ | Ref.SI.13 |
| Os-citrate<br>NPs | 0.035 | 12.15 | - | $H_2O_2$ | Ref.SI.14 |
| BN-GDY | 0.086 | 0.2058 | - | $H_2O_2$ | Ref.SI.15 |
| MCCP | 2.02 | 0.004 | 3.49 | $H_2O_2$ | Ref.SI.16 |

