## Supporting list for "Consolidated sustainable organic nanozyme integrated with Point-Of-Use sensing platform for dual agricultural and biological molecule detection"

Electronic Supplementary Information

**SI Figure 1.** Graphical illustration of the conceptualization of OM nanozyme and its expected enzyme-like catalytic pathway.
**SI Figure 2.** Additional TEM image of OM nanozyme.
**SI Figure 3.** Additional SEM image for reverse configuration.
**SI Figure 4.** Luminescence spectroscopical profile of OM nanozyme.
**SI Figure 5.** FT-IR analysis of the OM nanozyme.
**SI Figure 6.** Additional XPS quantification and deconvolution information.
**SI Figure7.** Supporting NTA (nanoparticle tracking analysis) profile of OM nanozyme.
**SI Figure 8.** Supporting EPR analysis of OM nanozyme.
**SI Figure 9.** Optimization and characterization of experimental condition of substrate-dependent colorimetric assay.
**SI Figure 10.** Storage stability of OM nanozyme a: temperature dependent comparison.
**SI Figure 11.** Schematic of the validation of the mixture selectivity of glyphosate.
**SI Figure 12.** Time dependent absorption compassion of the mixture selectivity of glucose.
**SI Figure 13.** Comparison of the glyphosate detection system with the conventional analytic tool.
**SI Figure 14.** Comparison of the glucose detection system with the conventional analytic tool.
**SI Figure 15.** Characterization of the paper-microfluidic chip using confocal microscopy.
**SI Figure 16.** Additional characterization of paper microfluidic chip
**SI Figure 17.** SEM analysis of OM nanozyme mounted on the paper-microfluidic chip.
**SI Figure 18.** Performance comparison between smartphone readout and the UV-VIS spectroscopy.
**SI Figure 19.** Linear graph of the $H_2O_2$ colorimetric assay/smartphone readout analysis on paper microfluidic chip.
**SI Figure 20.** Linear graph of the molecule detection/smartphone readout analysis on paper microfluidic chip.
**SI Figure 21.** Simplified(partially revised) Python code for obtaining the best $R^2$.
**SI Figure 22.** The formula for the fitting.
**SI Table 1. Kinetic profile table.**
